# Benchmarking of AlphaFold2 accuracy self-estimates as empirical quality measures and model ranking indicators and their comparison with independent model quality assessment programs

**DOI:** 10.1101/2023.12.15.571846

**Authors:** Nicholas S. Edmunds, Ahmet G. Genc, Liam J. McGuffin

## Abstract

**Motivation:** Despite an increase in the accuracy of predicted protein structures following the development of AlphaFold2, there remains a gap in the accuracy of predicted model quality assessment scores when compared to those generated with reference to experimental structures. The predictions of model accuracy scores generated by AlphaFold2, plDDT and pTM, have become familiar descriptors of model quality. However, at CASP15 some modelling groups noticed a variation in these scores for models of very similar observed quality, particularly for quaternary structures. There have also been a number of methods describing adaptations of the AlphaFold2 algorithm to purposes such as refinement by custom template recycling and model quality assessment using a similar method of template input. In this study we compare plDDT and pTM to their observed counterparts lDDT (including lDDT-Cα and lDDT-oligo) and TM-score to examine whether they retain their reliability across the whole scoring range for both tertiary and quaternary structures and in situations where the AlphaFold2 algorithm is adapted to customised functionality. In addition, we explore the accuracy with which plDDT and pTM rank AlphaFold2 tertiary and quaternary models and whether these can be improved by the independent model quality assessment programs ModFOLD9 and ModFOLDdock.

**Results:** For tertiary structures it was found that plDDT was an accurate descriptor of model quality when compared to observed lDDT-Cα scores (Pearson ρ = 0.97). Additionally, plDDT achieved a tertiary structure ranking agreement with observed scores of 0.34 as measured by true positive rate (TPR) and ModFOLD9 offered similar but not improved performance.

However, the accuracy of plDDT (Pearson ρ = 0.67) and pTM (Pearson ρ = 0.70) became more variable for quaternary structures quality assessment where overprediction was seen with both scores for models of lower quality and underprediction was also seen with pTM for models of higher quality. Importantly, ModFOLDdock was able to improve upon AF2-Multimer quaternary structure model ranking as measured by both TM-score (TPR 0.34) and lDDT-oligo (TPR 0.43). Finally, evidence is presented for an increase in variability of both plDDT and pTM when custom template recycling is used, and that this variation is more pronounced for quaternary structures.

## 1. INTRODUCTION

Since the success of AlphaFold2 (Jumper *et al*., 2021) at CASP14 in 2020 many articles have detailed the methodology by which AF2 achieved its level of accuracy, most notably by the DeepMind group itself (Evans *et al*., 2022) as well a group led by Jeffrey Skolnick (Jeffrey Skolnick *et al*., 2021) and the group who pioneered the development of the ColabFold adaptation of the software (Mirdita *et al*., 2022). It is usual for protein modelling software to provide accuracy self-estimate scores to accompany their models (Varadi *et al*., 2022) and while competition modellers are concerned with correlation agreements and statistical measures of significance across large datasets, the accuracy and usefulness of a single predicted score for one or only a few models may be more important to the general biologist. AlphaFold2’s state-of-the-art predicted models are increasingly relied upon and so it is vital that their accuracy is independently verified. In straightforward tertiary structure modelling AlphaFold2’s predicted lDDT score (plDDT) has been considered a useful indicator of quality (Yuma Takei and Ishida, 2022), but it is unclear whether this reliability transfers to quaternary structure modelling and whether there are any occasions on which the accuracy of these scores should be questioned.

### 1.1. AlphaFold2 predictions of model accuracy (plDDT, PAE and pTM)

plDDT is based on the local distance difference test (lDDT) (Mariani *et al*., 2013) which compares distances between individual atoms to estimate confidence in the arrangement of amino acid residues in the local environment. It is useful for assessing the local accuracy of domains, for example, as it will not penalise incorrect relative orientations of domains within a model of a multi-domain protein if there is a good match between the inter-atomic distance matrices. AF2 provides local plDDT per-residue scores in the B-factor column of a model’s coordinates file and a global per-model score which is output in the modelling log file.

The plDDT score itself is derived from the lDDT-Cα score (Tunyasuvunakool *et al*., 2021) which considers only the backbone Cα atoms in the distance calculation rather than the full all-atom lDDT score. It has a range of 0-100 (but lDDT values are also quoted as decimals in the 0-1 range), with high scores indicating higher confidence (Jumper *et al*., 2021). In general, plDDT values ≥ 90 equate to high confidence, those between 90 and 70 as confident, from 70 to 50 as low confidence and <50 as very low confidence with a tendency for disorder (Varadi *et al*., 2022). These confidence levels mean that plDDT scores are somewhat different to regular all-atom lDDT scores. Pfam (Stroe, 2021), for example, considers lDDT scores of ≥0.6 as representing reasonable models, 0.7 as good quality models and those above 0.8 as great models.

PAE represents the Predicted Aligned Error for residue backbone atoms, measured in Ångströms and calculated for each residue. Values are designed to measure the confidence in the predicted super-position of any two residues within the model and the native structure and it can be used to compare the residue confidence scores within a domain to those between domains. Lower scores represent low predicted error and therefore higher confidence, and higher scores (capped at 31.75) (Varadi *et al*., 2022) represent higher predicted error and therefore lower confidence. PAE is output as a colour-coded image mapping the areas of high and low confidence and also as machine-readable Json-formatted individual residue scores.

pTM is based on the topological similarity score TM-score (Zhang and Skolnick, 2004) and is calculated from the PAE matrix (Wallner, 2023). In later AlphaFold2 versions this is also output in the modelling log file and has a range of 0-1. No published confidence boundaries could be found for pTM but, traditionally, a TM-score of 1.0 would suggest a perfect match between a model and its native structure, a score greater than 0.5 is mostly interpreted as representing the same globular fold and scores below 0.17 are associated with unrelated proteins (Zhang and Skolnick, 2004). However, Jumper et al. (2021) described a relationship between pTM and TM-score as TM-score = 0.98 × pTM + 0.07 and so it may be appropriate to artificially construct pTM confidence boundaries using this relationship, if desired.

This study will concentrate on plDDT and pTM only for three simple reasons; PAE is not automatically normalised into an overall value meaning plDDT and pTM are the most often quoted AlphaFold2 confidence metrics; AF2 models are ranked by plDDT and AF2-Multimer models are ranked by pTM (Richard Evans, 2021) (see Supplementary section S1.1 for Evans’ description and ColabFold versions to which it applies), and that these scores are familiar and directly measurable against their observed counterparts, lDDT and TM-score.

### 1.2. Documented descriptions of AlphaFold2 predicted scores

One of the strengths of the AF2 algorithm has been described as its ability to recognise low accuracy local areas (Shao *et al*., 2022) or indeed whole models and apply confidence scores appropriately. As stated above, linear relationships have been described (Jumper *et al*., 2021) for lDDT-Cα as 0.997 × plDDT − 1.17 and TM-score as 0.98 × pTM + 0.07. While these relationships acknowledge a tendency for some over-prediction with plDDT, the suggestion is that both scores are consistently applied across the scoring range. However, at CASP15 (2022), despite plDDT and pTM scores successfully contributing to many groups’ model-selection algorithms, it was noticed that there was a variability in these scores, particularly for multimer models of very similar quality. One group (Wallner, 2023) reported that up to one-third of models with a ranking confidence of pTM > 0.8 actually had the wrong domain orientation and our own experiences during CASP15 modelling revealed an increase in plDDT as high as 40 points during model refinement, which would suggest an overprediction of model quality improvement.

### 1.3. Wider uses of AlphaFold2 rely on accurate predicted quality

Since the CASP14 success detailed above, AlphaFold2 has been used in a DeepMind-EMBL collaboration to create the AlphaFold Protein Structure Database https://alphafold.ebi.ac.uk (Tunyasuvunakool *et al*., 2021). This is aimed at creating a community resource allowing easy access to protein structures which remain unsolved by traditional experimental methods. With the growing profile of in-silico modelling against the backdrop of a growing community investment in artificial intelligence (AI), databases such as this are likely to increase in popularity along with a greater reliance on computational modelling software. Although, for now, the database is limited to tertiary structures, it might, nevertheless, be prudent to examine whether AlphaFold2’s confidence metrics can be relied upon to rate and rank models accurately across the whole quality range. Further to this, at least three published works describe using the AlphaFold derivative ColabFold to input models as custom templates. One group (Terwilliger, 2022) input electron density maps during model generation from experimental data, another (Adiyaman *et al*., 2023) described a procedure for model improvement using custom template recycling as a refinement strategy, and a third (Roney and Ovchinnikov, 2022) described a method for using AlphaFold2 as a quality assessment tool.

It is clear, then, that a certain reliance is being placed on plDDT and pTM scores and this study aims to assess the performance of these scores in both monomer and multimer model populations in comparison to their observed lDDT and TM-score counterparts. Within the model populations, models will be generated both with and without custom template recycling to evaluate whether there is a variation in predictive performance with this single variable. In addition plDDT and pTM will be compared to quality scores generated by the independent quality assessment programs ModFOLD9 (tertiary structures) and ModFOLDdock (quaternary structures) (McGuffin *et al*., 2023).

### 1.4. Objectives

Using, blind modelling and assessment data from CASP15, the relationship between AlphaFold2 predicted scores and their observed counterparts, lDDT score (including lDDT-Cα and lDDT-oligo) and TM-score, will be examined in four parts.

First, we will assess the scores’ accuracy at describing tertiary and quaternary structures in terms of both global model quality and ranking agreement with observed scores. Two hypotheses have been formulated to test this.

1. *H0. There is no increase in magnitude between the AF2 predicted and equivalent observed scores. H1. The magnitude of the AF2 predicted scores is higher than the equivalent observed scores*.
2. *H0. There is no association between the AF2 predicted and observed score ranking categories. H1. There is an association between the AF2 predicted and observed score ranking categories*.

Second, blind prediction scores used for the CASP15 EMA competition will be used to examine the comparative performance between ModFOLDdock and AF2-Multimer scores for quaternary structures. Similarly, ModFOLD9 predictions, which were also blind and run in house prior to the release of the CASP15 experimental structures, will be used to examine the performance between AlphaFold2 and ModFOLD9 scores for tertiary structures.

3. *H0. There is no difference between the independent QA and AF2 rankings as measured by the association between model rank categories. H1. Independent QA and observed score model ranks are more closely associated than AF2 and observed score model ranks*.

Finally, we examine the effect of custom template recycling on the accuracy of the AF2 and AF2-Multimer predicted scores. These results are described in Supplementary section S3.4.

4. *H0. There is no difference between AF2 regular modelling and custom template modelling predicted scores, when compared to equivalent observed scores. H1. AF2 predicted scores following custom template modelling show greater variation than scores from regular modelling, when compared to equivalent observed scores*.

## 2. Materials and Methods

### 2.1. Selection of models to test the hypotheses

Four individual datasets were used for this study.

Population A (CASP15 monomers) comprised the McGuffin group’s tertiary structure submissions for CASP15. Population B (CASP15 multimers) was composed of both the McGuffin group’s (MultiFOLD, group 462) and the ColabFold group’s (group 446) multimer submissions for CASP15. Group 446 submissions are publicly available from https://casp15.colabfold.com/). Population C (recycled monomers) is a superset of the AF2 and non-AF2 models used in the custom-template recycling experiment described in our recycle paper (Adiyaman et al., 2023). The original model population was fixed at 16 CASP14 targets to form a common subset with the ReFOLD4 molecular dynamics analysis which used only the FM targets submitted by the AlphaFold group. The emphasis of this experiment has shifted from measuring model improvement to measuring model quality overprediction and so it was felt that the inclusion of four additional FM/TBM targets, for which scores had already been collected, was justified to increase the model population without significantly altering the difficulty of the models. This increased the total target number to 20. Population D (recycled multimers) is the same multimer population used in the custom-template recycling experiment described in our recycle paper.

### 2.2. The Population A dataset – CASP15 monomers

This consisted of all McGuffin group’s blind model submissions for 26 CASP15 regular tertiary structure targets for which ModFOLD9 scores and a reference native structure were available.

Our group’s modelling algorithm used two separate rounds. Round 1 used regular modelling only, with no refinement process, whereas round 2 included refinement by custom template recycling. Models were therefore split int two sub-populations; Population A1 represented the round 1 models (regular modelling) and these were created with a default of 12 recycles and both with and without AMBER relaxation, resulting in 20 models per target (5 unrelaxed AF2, 5 relaxed AF2, 5 unrelaxed AF2M and 5 relaxed AF2M). For a small minority of large targets memory constraints meant relaxation was not always possible resulting in fewer models. Population A2 represented the round 2 models which were subject to custom template recycling and resulted in 10 models per target. Again, 5 of these underwent AMBER relaxation while the other 5 remained unrelaxed. In this way a maximum of 30 models were created per target. Predicted plDDT and pTM scores were harvested directly from the server for both sub populations and predicted ModFOLD9 scores were collected from the original cached datasets used during CASP15. Observed lDDT and TM-scores were generated using the downloadable versions of TM-score (Zhang and Skolnick, 2004) and lDDT score (Mariani *et al*., 2013) to compare models for each target with the CASP observed structures. A total of 735 models were analysed; consisting of 490 round 1 and 245 round 2 models.

### 2.3. The Population B dataset – CASP15 multimers

This model population comprised all blind multimer (assembly) CASP15 model submissions for both the MultiFOLD (462) and ColabFold (446) group servers. These two sets of models were chosen because they were created using the same base ColabFold software (although exact versions may differ slightly) but differed by the use of custom template recycling in the MultiFOLD pathway. The rationale was that the ColabFold models could be used to assess AF2-Multimer score overprediction when only regular modelling was used and, that by comparing the ColabFold and MultiFOLD populations, the effect of the additional custom template recycling on predicted scores could be assessed.

The ColabFold group multimers are named Population B1. For these, custom template recycling and AMBER relaxation were not used and 12 recycles was used as default (Sergey Ovchinnikov *et al*., 2022). The predicted scores, plDDT, pTM (and iPTM where available) were harvested directly from the server website. The MultiFOLD group models are named Population B2. The same pathway as outlined in 2.2 above, including custom template recycling, was used to create these models. Only the final 5 models submitted to CASP were used for analysis and again predicted scores were collected directly from the server. For comparisons with observed scores, the official CASP15 assessor lDDT-oligo and TM-scores were downloaded from the CASP15 prediction centre results webpage. As the ModFOLDdock server participated in the CASP15 EMA experiment, predicted ModFOLDdock and ModFOLDdockR scores were also readily available for both sub populations of models. Scores for rank 1 to 5 models were collected for all multimer models for which data were available, resulting in 395 individual models across 41 targets (the ColabFold group submitted no models for three targets making a total of 38). In total the Population B dataset consisted of 395 multimer scores. The CASP targets used in populations A and B are listed in Supplementary section S2.2 and S2.3.

### 2.4. The Population C dataset – recycled monomers

This dataset consisted of the custom template recycled AlphaFold2 and non-AlphaFold2 tertiary models with the addition of four extra targets as explained in section 2.1 above. The processing of models can be found in Supplementary section S2.4. The AF2 dataset contributed 800 individual scores from 8 sets of scores per model across 5 models per target for 20 targets. Non-AF2 models were selected from the same 20 FM targets for the next five best-ranked groups beneath AlphaFold2 at CASP14. These were Baker (473), Baker-experimental (403), Feig-R2 (480), Zhang (129) and tFold_human (009). To ensure consistency in terms of globular fold similarity, only models with a TM-score >=0.45 were selected and this resulted in 47 individual models with a total of 1880 individual model scores.

### 2.5. The Population D dataset – recycled multimer models

This dataset consisted of custom template recycled multimer models. As the AlphaFold2 group did not submit multimer (assembly) models at CASP14, models for this dataset were selected from the CASP14 top five ranked groups. According to official results tables, these were Baker, Venclovas, Takeda-Shitaka, Seok and DATE. See Supplementary section S2.5 for a description of the model processing. This dataset contributed a total of 2000 individual scores. An overall total of 5,810 model scores were collected across the whole study.

The method for handling multi-contingency table data and ranking by pTM is also listed in Supplementary section S2.6.

## 3. Results and Discussion

### 3.1 Hypothesis 1

#### Are AF2-predicted scores higher than the equivalent observed scores?

In order to focus on one independent variable at a time, the question of whether predicted scores are good quality indicators must be answered using only models which have *not* undergone custom template recycling. For monomers, this is population A1 (round 1 models) and for multimers this is population B1 (ColabFold multimers). Population A1 will be considered first.

#### 3.1.1. Part 1. Monomer data; Population A1, (round 1)

AF2 default monomer ranking is by plDDT and so results will focus on plDDT/lDDT similarity.

The plots in Figure 2 show that plDDT scores are slightly elevated compared to the all-atom lDDT scores. However, when plDDT scores are considered with reference to lDDT-Cα scores ((Jumper *et al*., 2021), (Tunyasuvunakool *et al*., 2021)) in Figure 3, there is no evidence of plDDT over-prediction, in-fact the boxplot in Figure 3 shows a slightly lower median score for plDDT. It should also be possible to check whether the plDDT values in this sample are in line with the published linear relationship described in section 1.2 (lDDT-Cα=0.997×plDDT− 1.17). If the median plDDT value of 0.91 from the Figure 3 boxplot is considered as a convenient example, the median lDDT-Cα score can be calculated from plDDT in three simple steps:

**Figure 2.**
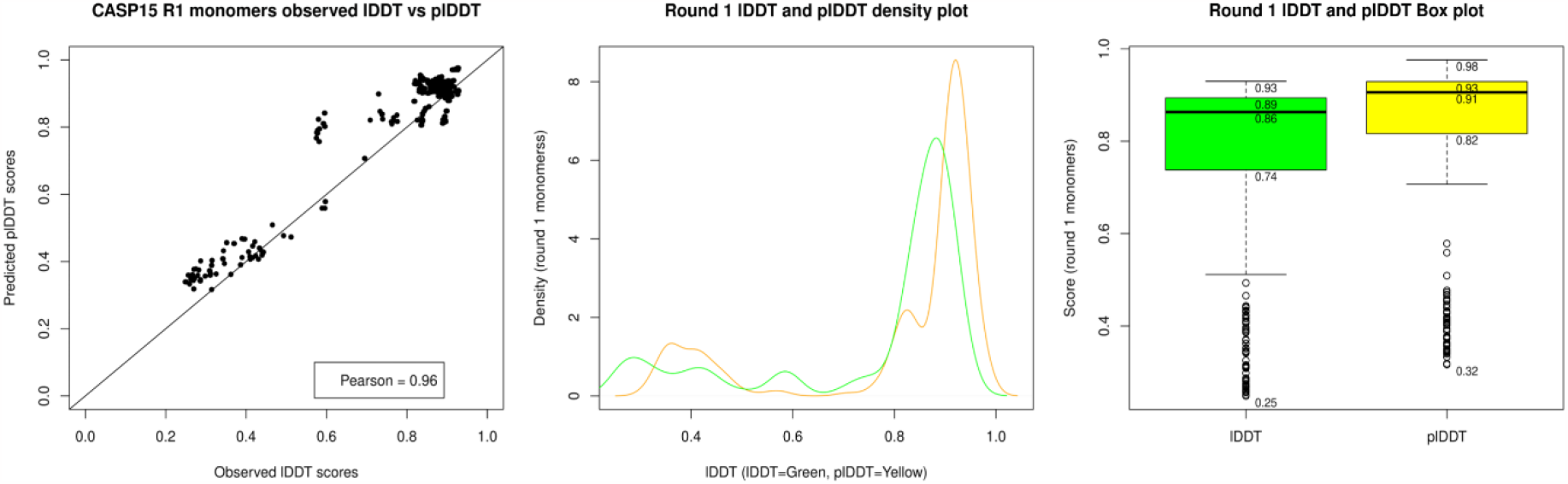
Plots of plDDT versus observed lDDT for round 1 monomers in population A1. A scatter plot (left), a density plot (middle) and a boxplot (right) for the same population. For all plots plDDT has been rescaled to fit the 0-1 lDDT range.

**Figure 3.**
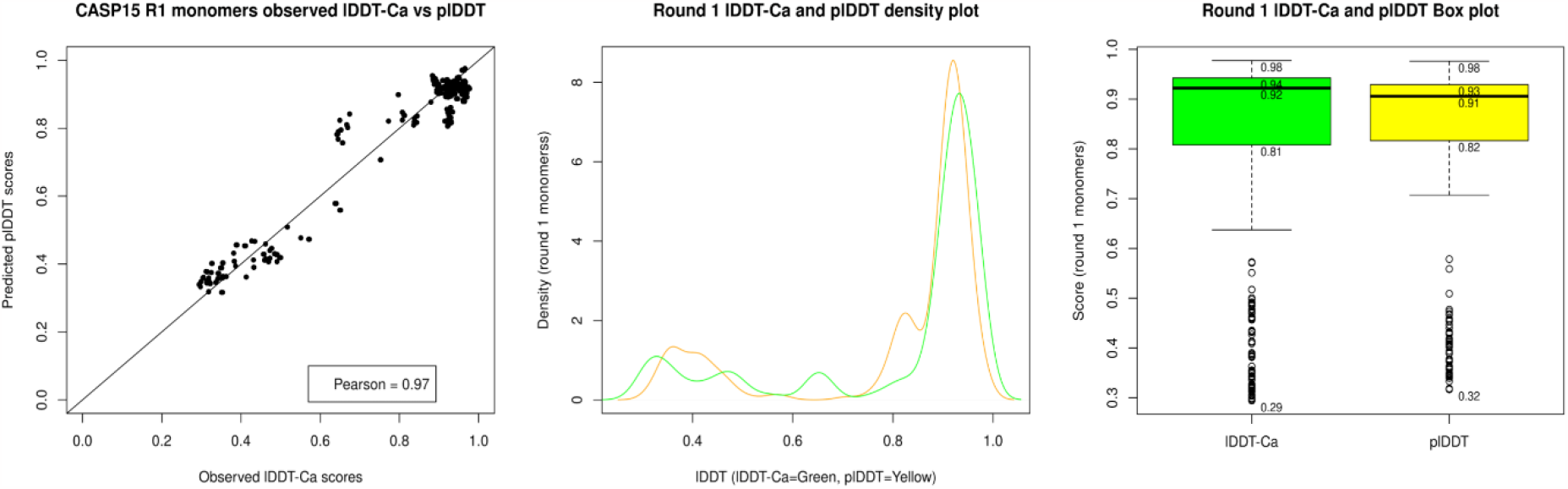
Plots of plDDT versus observed lDDT-Cα for round 1 monomers population A1. A scatter plot (left), a density plot (middle) and a boxplot (right) for the same population. Again, plDDT has been rescaled to the 0-1 range.

1. Convert plDDT back to its 0-100 range: 0.91 x 100 = 91.0
2. Calculate lDDT-Cα from the relationship: 0.997 x 91 - 1.17 = 89.56
3. Convert lDDT-Cα back to the 0-1 range: 89.56/100 = 0.8956 or 0.90 to 2.d.p.

From Figure 3 it can be seen that the actual lDDT-Cα is 0.92, meaning that rather than being overpredicted, plDDT has in fact been slightly underpredicted for this sample of models.

To formally test this data against hypothesis 1, a Wilcoxon signed rank test for non-parametric paired data was carried out to test significance. The following results were obtained (a Shapiro test for normality gave p-values of <0.05 for all three (plDDT, lDDT and lDDT-Cα) scores, showing the distributions to be non-normal in all cases).

Table 2 results show that, for this sample of monomers, there is a significant difference between predicted and observed lDDT scores with P-values of 2.2×10^−16^ and 9.69×10^−6^ for the 2-sided Wilcoxon tests for all atom lDDT and lDDT-Cα respectively. However, there is disagreement between the two scores, with row 2 of the table showing that according to a 1-sided test, plDDT values are greater than those for all atom lDDT (p-value of 2.2×10^−16^) while row 4 shows the opposite, that lDDT-Cα values are actually significantly higher than plDDT values (p-value of 4.81×10^−6^). Considering the published works cited earlier confirming that plDDT is based on lDDT-Cα it would be more appropriate to accept the null hypothesis in this case. Therefore, for monomers constructed from regular straight-forward AF2 modelling and assessed by lDDT-Cα: *There is no increase in magnitude between the AF2 predicted and equivalent observed scores*.

**Table 1.** A summary of the different model populations used in the study.

| Model population | Type and modelling software | Stoichiometry and type of modelling |
| --- | --- | --- |
| Population A1 | CASP15, MultiFOLD round 1 | Monomer, regular modelling. |
| Population A2 | CASP15, MultiFOLD round 2 | Monomer, custom recycling included. |
| Population B1 | CASP15, ColabFold | Multimer, regular modelling. |
| Population B2 | CASP15, MultiFOLD | Multimer, custom recycling included. |
| Population C | CASP14, AF2 and non-AF2 | Monomer, custom recycling included. |
| Population D | CASP14, top 5 groups. | Multimer, custom recycling included. |

**Table 2.** Wilcoxon statistics for population A1, round 1 monomers.

| *Scores compared* | *Independence and distribution symmetry* | *p-value* |
| --- | --- | --- |
| *pIDDT versus IDDT* | Paired; 2-sided test. | **$2.2 \times 10^{-16}$** |
| *pIDDT versus IDDT* | Paired; 1-sided (pIDDT > IDDT) | **$2.2 \times 10^{-16}$** |
| *pIDDT versus IDDT-Cα* | Paired; 2-sided test. | **$9.69 \times 10^{-6}$** |
| *pIDDT versus IDDT-Cα* | Paired, 1-sided (pIDDT < IDDT-Cα) | **$4.81 \times 10^{-6}$** |

#### 3.1.2 Part 2. Multimer data; Population B1 (ColabFold multimers)

For multimers pTM is the default ranking metric, however plDDT was used in early versions of ColabFold and so both scores are considered here.

Both the scatter and density plots in Figure 4 appear to show an under-estimation of pTM score for higher quality multimer models but a relatively large overestimation for some lower-quality models. For Figure 5, plDDT appears to be over-estimated across the quality range which may be accounted for by the use of an all-atom observed lDDT-oligo score. However, as with pTM scores, there is a more pronounced overestimation for some models in the lower quality range. The Shapiro test for normality (all scores were non-normal) and Wilcoxon signed rank test for significance were executed in the same way as described for monomer data.

**Figure 4.**
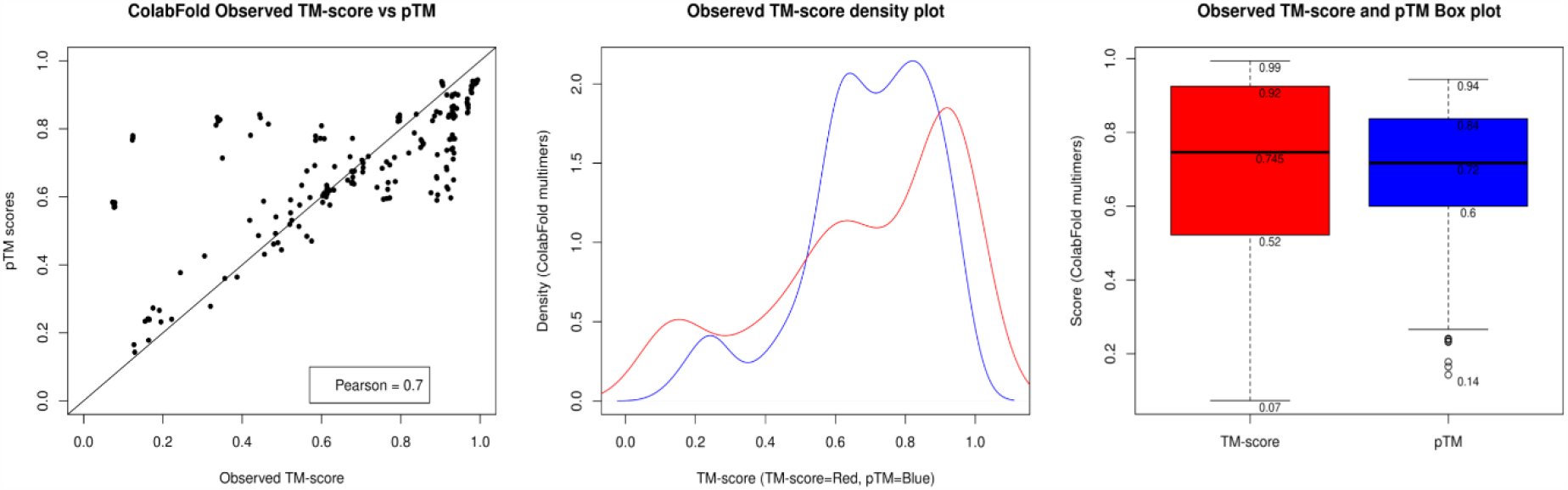
Plots for pTM versus observed TM-score for Population B1 (ColabFold multimers). A scatter plot (left), density plot (middle) and boxplot (right).

**Figure 5.**
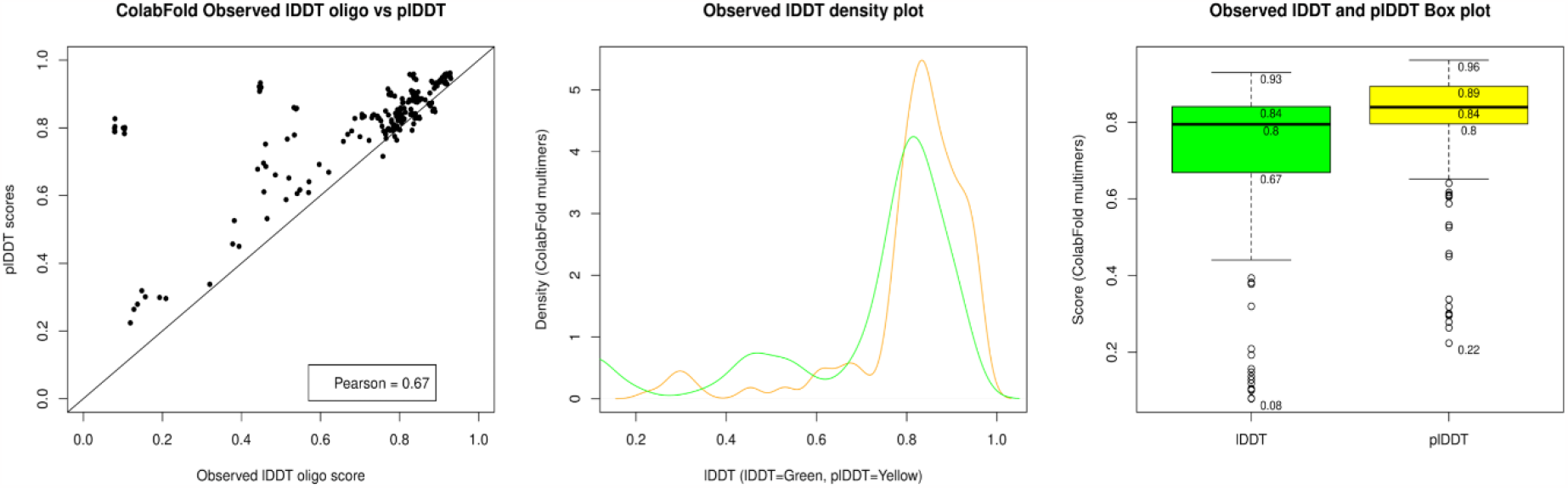
Plots for plDDT versus observed CASP lDDT-oligo for Population B1 (ColabFold multimers). A scatter plot (left), density plot (middle) and boxplot (right).

Table 3 shows that there is a significant difference between predicted plDDT and observed lDDT scores and that plDDT values are significantly higher than lDDT-oligo as shown by the p-value of 2.2×10^−16^. For hypothesis 1, with respect to lDDT, the alternative hypothesis can therefore be accepted for ColabFold multimers, i.e., *The magnitude of the AF2 predicted scores is higher than the equivalent observed scores*.

**Table 3.** Wilcoxon statistics for population B1, ColabFold multimers. *(significant figures in bold)*

| **Scores compared** | **Independence and distribution symmetry** | **p-value** |
| --- | --- | --- |
| *pIDDT versus IDDT-oligo* | Paired; 1-sided test, pIDDT > IDDT-oligo | **$2.2 \times 10^{-16}$** |
| *pTM versus TM-score* | Paired, 2-sided | **0.038** |
| *pTM versus TM-score* | Paired; 1-sided test, pTM > TM-score | 0.980 |
| *pTM versus TM-score* | Paired; 1-sided test, pTM < TM-score | 0.0192 |

**Table 4.** The summary statistics for round 1 monomer ranking agreement.

| Test | IDDT result | IDDT-Ca result |
| --- | --- | --- |
| Macro-Sensitivity (TPR) | 0.3204 | 0.3428 |
| Macro-Specificity | 0.8301 | 0.8357 |
| Macro-Precision | 0.3204 | 0.3428 |
| Macro-Accuracy | 0.7281 | 0.7371 |
| Fisher's Exact (p-value) | < 0.001 | < 0.001 |
| Chi-squared ($\chi^2$; p-value) | 128.27; $2.2 \times 10^{-16}$ | 167.35; $2.2 \times 10^{-16}$ |

The data are not so clear for TM scores. There is a significant difference between pTM and TM-score but rather than pTM being the greater of the two (p-value of 0.980), TM-score may in fact be greater than pTM (p-value of 0.019). To reveal more information about the relationship between pTM and TM-score, a further investigation into the variation in the two scores is described in Figure 6 below.

**Figure 6.**
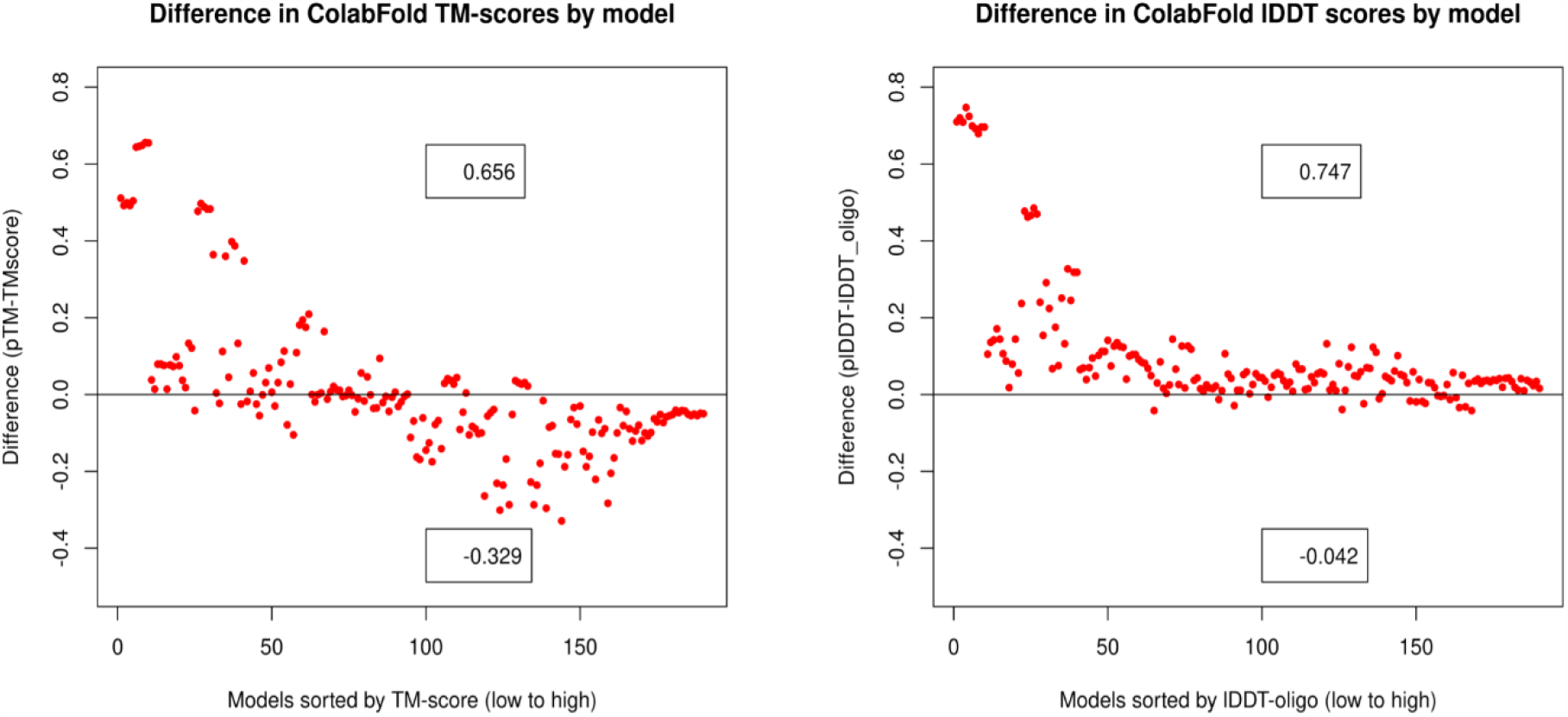
Two plots showing the difference between predicted and observed scores for population B1 (ColabFold multimers). The line at 0.0 represents the observed score; predicted scores are represented as points. Left: pTM versus TM-score, right: plDDT versus lDDT-oligo score. Numbers on the x-axis are the models in the population, ordered from low to high observed score.

The relationships suggested in Figure 5 and Table 3 are more clearly shown by the two plots in Figure 6. Both plots show that an overestimation of predicted scores is more likely for lower quality models with a maximum difference of +0.65 for pTM and +0.74 for plDDT. Again, a tendency for underestimation of pTM in higher quality models is apparent with a maximum difference of −0.32. This explains the Wilcoxon test results for pTM; there is both over and under-estimation occurring which is quality-related and which, to some extent, cancel each other out. While there is an allusion to minor pTM underprediction in the mathematical relationships described in section 1.2 (Jumper *et al*., 2021), no documentation relating to an overprediction for lower quality models could be found. A similar pattern of underestimation is not seen for plDDT.

For hypothesis 1, with respect to TM-score, the null hypothesis must be accepted for ColabFold multimers, i.e., *There is no increase in magnitude between the AF2 predicted and equivalent observed scores*. However, a caveat must be added to this last statement, that, for this population (intended to represent regular multimer modelling), while a significant increase in predicted TM-score could not be detected in the overall population, overprediction was observed in models of lower observed quality.

### 3.2 Hypothesis 2

#### Is AlphaFold2 model ranking reliable compared to ranking by observed scores, as measured by association between model rank categories?

Again, to answer this question fairly, models which have not undergone custom template recycling must be used. Therefore, this analysis will use the same data as used in 3.1 - Population A1 (round 1 models) and Population B1 (ColabFold multimers).

The contingency tables in Figure 7 show strong agreement for monomer data between observed lDDT-derived ranks and plDDT predicted ranks with a slightly stronger agreement when lDDT-Cα is used as the observed measure. The level of agreement for rank 1 and rank 5 data shown in the contingency tables is supported by mean true positive rates (TPR) of 32.04% and 34.28% for lDDT and lDDT-Cα respectively. In addition, the Fisher’s exact tests return p-values well below the significance level of 0.05 and the Chi-squared tests return values of 128.27 (lDDT) and 167.35 (lDDT-Cα) with very small p-values. These data provide robust evidence that this distribution was unlikely to occur by chance and that there is a significant positive relationship between the predicted and observed scores.

**Figure 7.**
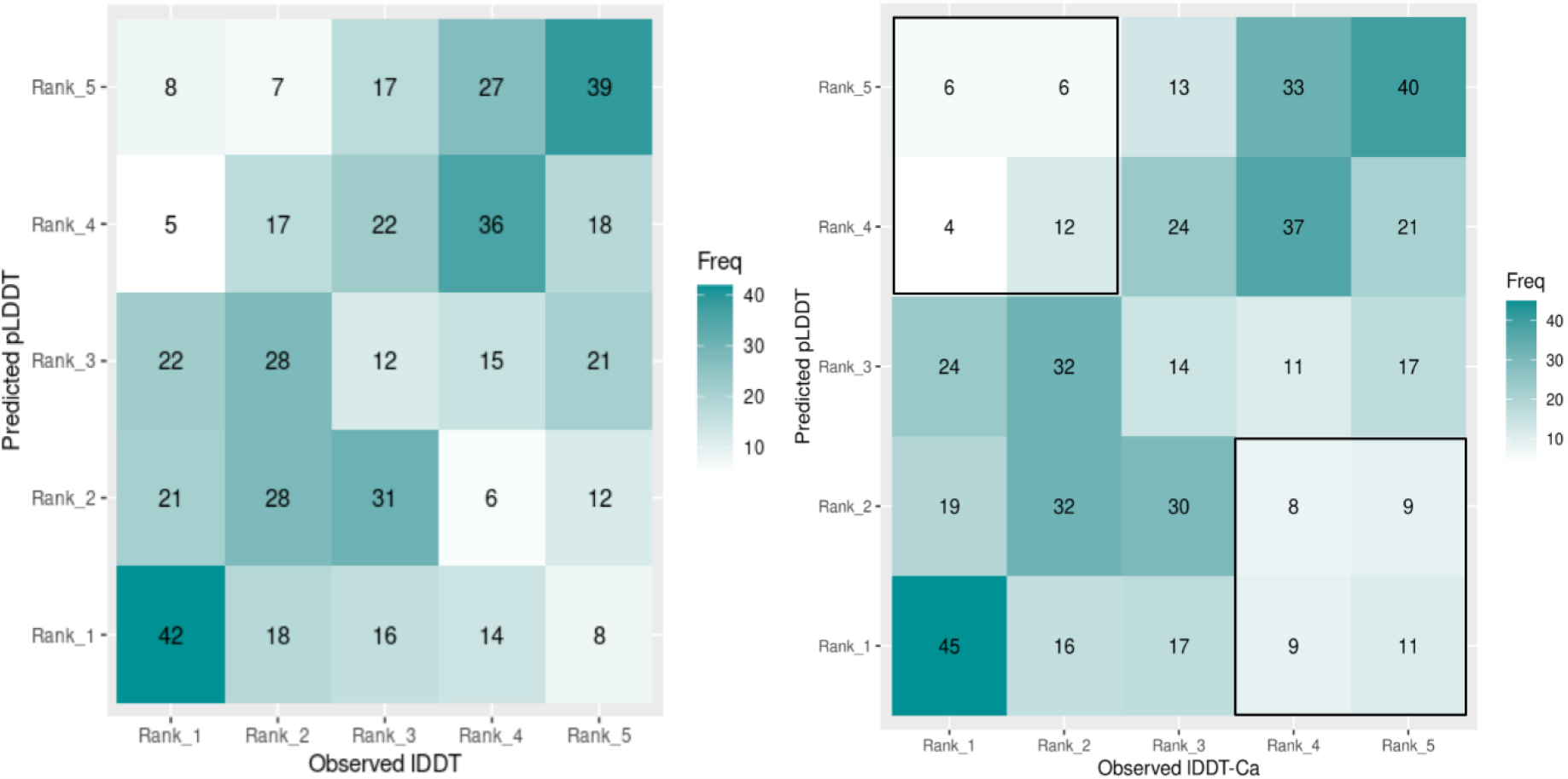
Contingency tables showing the rank agreement between observed lDDT and plDDT values for Population A1 (round 1 monomers). Left, for all-atom observed lDDT scores; Right, for observed lDDT-Cα scores. Accompanying table of calculated statistics below.

For the multimer population represented by Figure 8, the agreement looks appreciably less certain for both pTM and plDDT scores. The summary statistics show a reduction in mean TPR to 30.5% for pTM and 28.4% for plDDT. Both Fisher’s exact and Chi-squared p-values, however, remain significant suggesting a relationship between the two rank sets, although it is notable that the magnitude of the χ^*2*^ statistic has decreased for both pTM and plDDT suggesting a weaker association between predicted and observed ranks.

**Figure 8.**
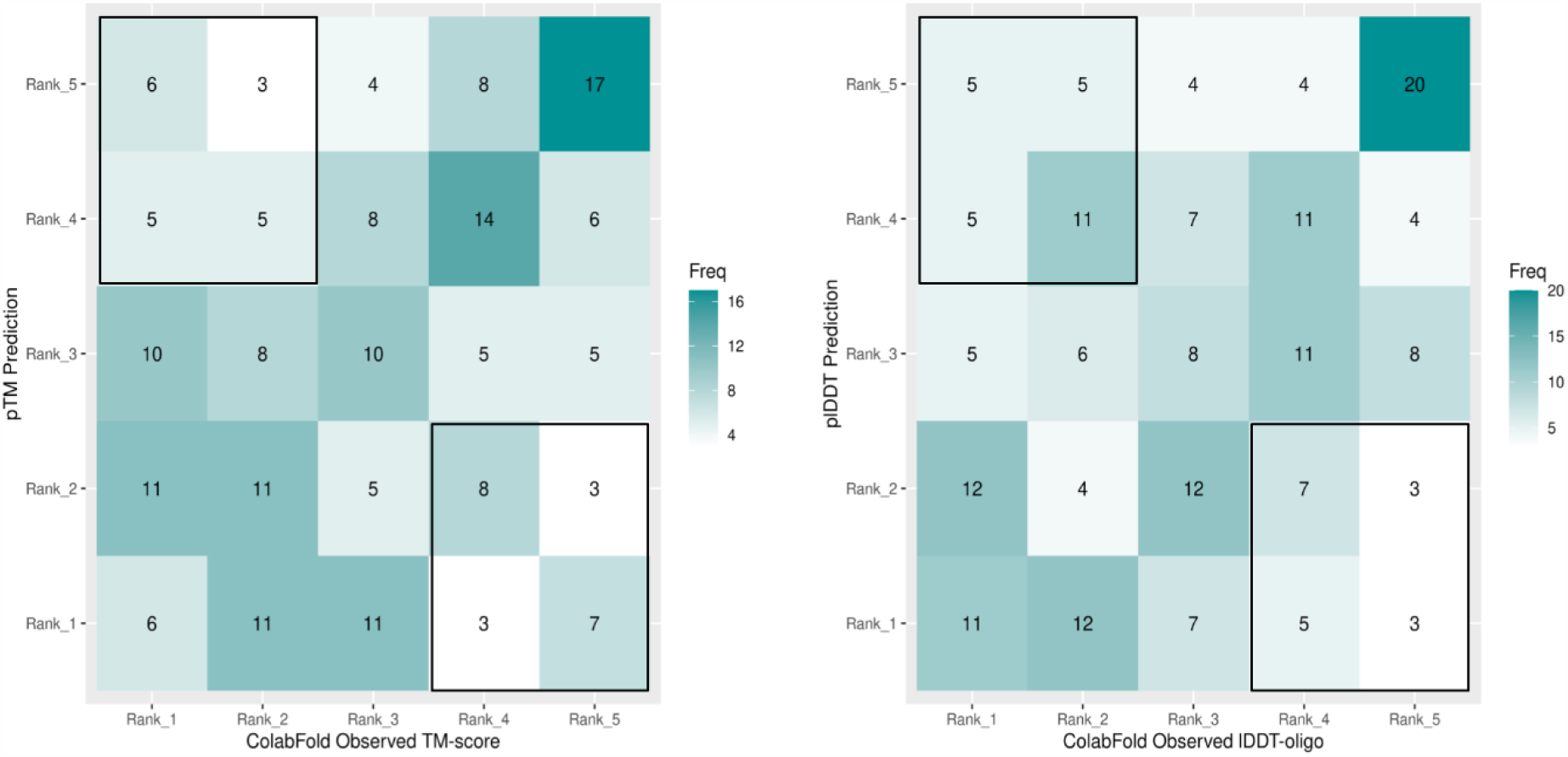
Contingency tables showing rank agreement for Multimers in Population B1 (ColabFold multimers). Left, observed TM-scores versus pTM and right, observed lDDT-oligo versus plDDT scores. Accompanying table of calculated statistics below.

For hypothesis 2, these results suggest that there is significant association between the distribution of predicted and observed ranks for both monomer and multimer model populations created via regular modelling and the alternative hypothesis can be accepted, i.e., *There is an association between the AF2 predicted and observed score ranking categories*. Similarly, to above, though, a qualifying statement may be appropriate here to add that the association appears more robust for tertiary structure ranking by plDDT than for multimer ranking by either plDDT or pTM.

### 3.3 Hypothesis 3

#### Can model ranking accuracy be improved by independent MQA programs?

The individual rank-agreement and TPR values described above for monomer and multimer models need to be contextualised by comparison to another leading QA method. This section presents identical analysis for ranking based on predicted scores from the independent QA programs ModFOLD9 (monomer data) and ModFOLDdock (multimer data).

Visual comparison of the data in Figure 9 to those in Figure 7 shows that ModFOLD9 has been unable to improve upon the ranking agreement between plDDT and lDDT scores for monomers. TPR is reduced from 34.2% to 26.9% (lDDT-Cα) and all other macro-averaged statistics are lower than previously reported. Although the Fisher’s exact and Chi-squared tests continue to return significant p-values, the χ^*2*^ values, in agreement the TPR scores, have reduced suggesting a weaker overall association between the ranks.

**Figure 9.**
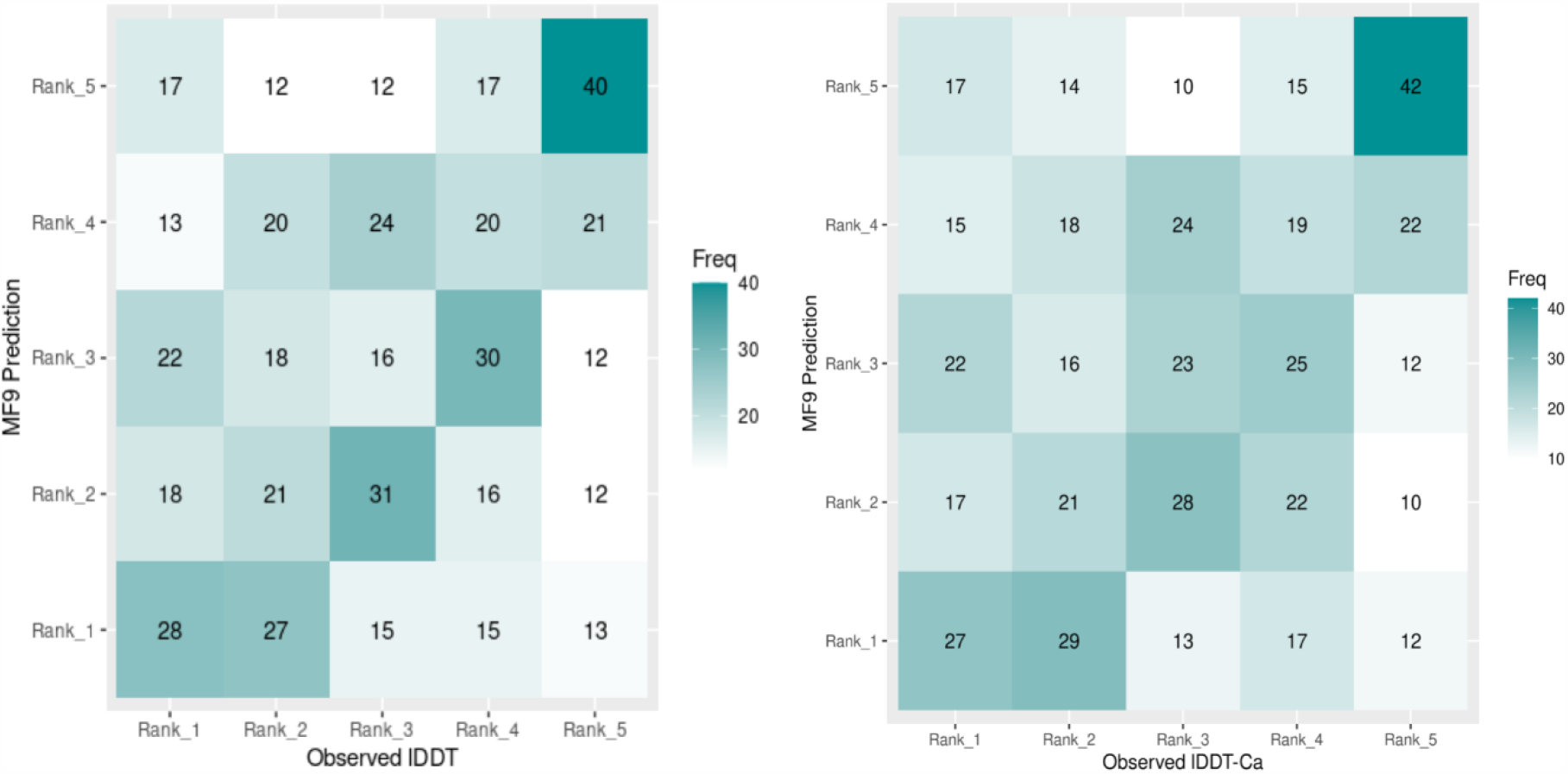
Contingency tables showing the rank agreement between observed lDDT and ModFOLD9 values for Population A1 (round 1 monomers). Left, using all-atom lDDT scores; right, using observed lDDT-Cα scores. Accompanying table of calculated statistics below.

Therefore, the closeness of the relationship has not been improved by ModFOLD9 and for hypothesis 3, in respect to ModFOLD9, the null hypotheses must be accepted, i.e., *There is no difference between the independent QA and AF2 rankings as measured by the association between model rank categories*.

In contrast, a visual comparison of the data in Figure 10 with those from Figure 8 suggests ranking agreement for multimers is stronger for ModFOLDdock scores, particularly for lDDT rank agreement. This is supported by the data in Table 7 where the TPR has increased from 30.5% in Table 5 to 34.2% in Table 7 for TM-score and more appreciably from 28.4% to 43.1% for lDDT-oligo score. The Chi squared values have remained similar for TM-score across the two tables, however Table 7 shows an increase in the χ^*2*^ statistic from 51.31 to 78.94 for lDDT-oligo ranking. This increase, along with the increased TPR values, is strongly suggestive of a closer positive association between ModFOLDdock predicted and lDDT-oligo ranking.

**Table 5.**
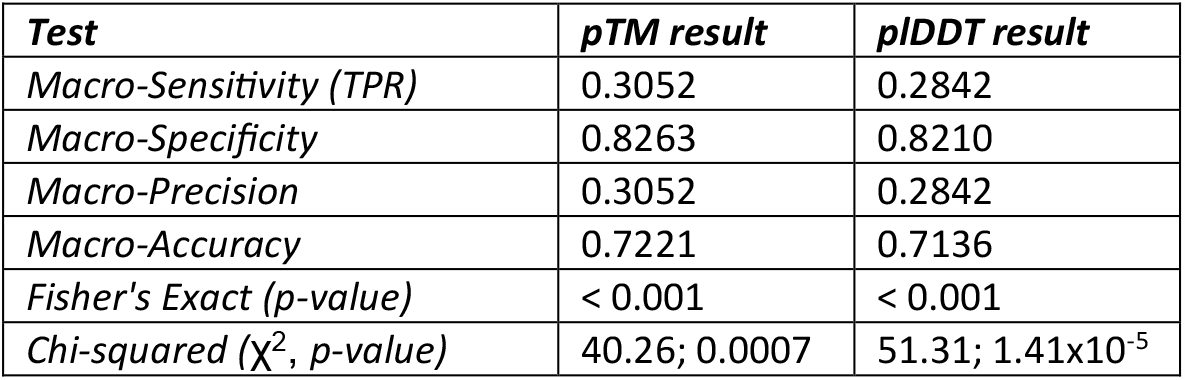
Summary statistics for ColabFold multimer ranking agreement.

| **Test** | **pTM result** | **pIDDT result** |
| --- | --- | --- |
| *Macro-Sensitivity (TPR)* | 0.3052 | 0.2842 |
| *Macro-Specificity* | 0.8263 | 0.8210 |
| *Macro-Precision* | 0.3052 | 0.2842 |
| *Macro-Accuracy* | 0.7221 | 0.7136 |
| *Fisher's Exact (p-value)* | < 0.001 | < 0.001 |
| *Chi-squared ($\chi^2$, p-value)* | 40.26; 0.0007 | 51.31; 1.41x10<sup>-5</sup> |

**Table 6.**
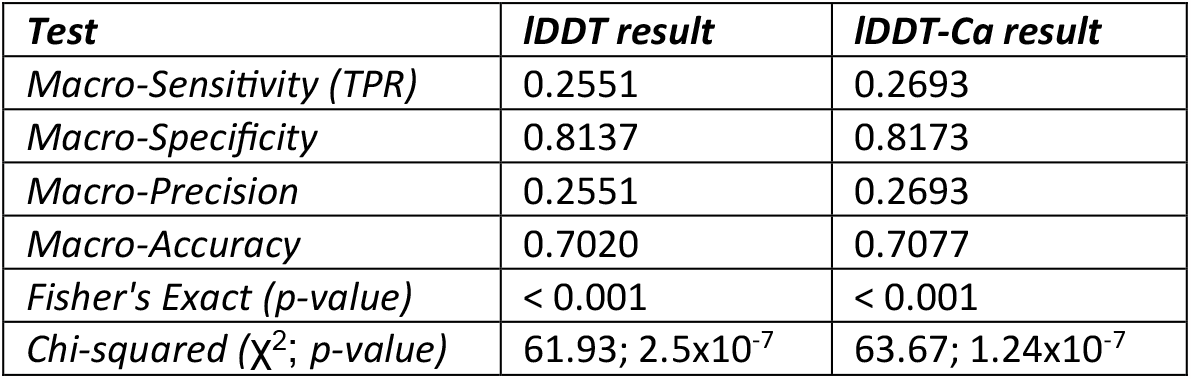
The summary statistics for round 1 monomer ranking agreement.

| Test | IDDT result | IDDT-Cα result |
| --- | --- | --- |
| Macro-Sensitivity (TPR) | 0.2551 | 0.2693 |
| Macro-Specificity | 0.8137 | 0.8173 |
| Macro-Precision | 0.2551 | 0.2693 |
| Macro-Accuracy | 0.7020 | 0.7077 |
| Fisher's Exact (p-value) | < 0.001 | < 0.001 |
| Chi-squared ($\chi^2$; p-value) | 61.93; $2.5 \times 10^{-7}$ | 63.67; $1.24 \times 10^{-7}$ |

**Table 7.** Summary statistics for ColabFold multimer ranking agreement.

| Test | TM-score Result | IDDT result |
| --- | --- | --- |
| Macro-Sensitivity (TPR) | 0.3421 | 0.4315 |
| Macro-Specificity | 0.8355 | 0.8578 |
| Macro-Precision | 0.3421 | 0.4315 |
| Macro-Accuracy | 0.7368 | 0.7726 |
| Fisher's Exact (p-value) | < 0.001 | < 0.001 |
| X-squared ($\chi^2$; p-value) | 38.42; 0.0013 | 78.94; 2.57x10<sup>-10</sup> |

**Figure 10.**
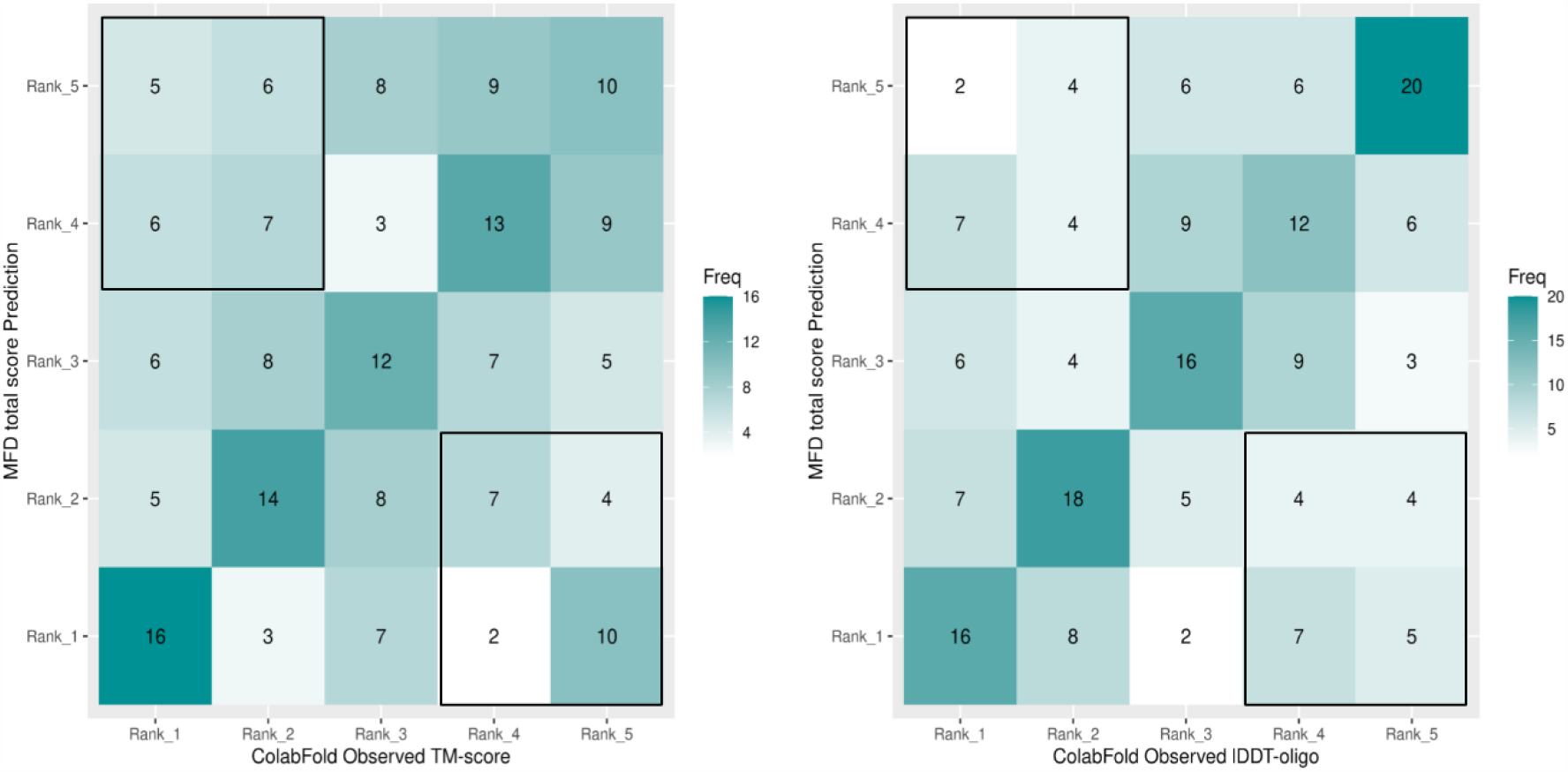
Contingency tables showing rank agreement for Population B1 (ColabFold multimers). Left, between observed TM-scores and ModFOLDdock score; right, between observed lDDT-oligo and ModFOLDdock score. Accompanying table of calculated statistics below.

For hypothesis 3, then, with respect to multimer ranking by TM-score, there is insufficient evidence to reject the null hypothesis. *There is no difference between the independent QA and AF2 rankings as measured by the association between model rank categories*.

However, with respect to multimer ranking by lDDT, considering the increases in scores described above, there may be sufficient evidence to accept the alternative hypothesis, i.e., *Independent QA and observed score model ranks are more closely associated than AF2 and observed score model ranks*.

## 4. Conclusions

Throughout, this study has been orientated toward answering four questions concerning the accuracy of the often-quoted AlphaFold2 predicted scores plDDT and pTM, both as empirical descriptors of model quality and as reliable ranking scores.

### plDDT is a reliable indicator of tertiary structure (monomer) model quality and ranking

From the data presented in section 3.1.1, plDDT was shown to be a reliable indicator of tertiary structure model quality when straight forward regular modelling was used and showed impressive Pearson correlation coefficients with both observed lDDT and lDDT-Cα scores which the independent QA method, ModFOLD9, was unable to improve upon. plDDT prediction accuracy appeared to be maintained across the scoring range and any over-prediction may be potentially explained by the published linear relationship with lDDT-Cα.

Ranking of the same tertiary model population also showed an agreement between plDDT and lDDT-Cα assigned ranks, which ModFOLD9 was again unable to improve upon. For straight forward regular tertiary modelling and it can be concluded that plDDT appears to be a reliable descriptor of quality and ranking tool for tertiary structure models.

### Both pTM and plDDT show variability as indicators of multimer model quality and ranking and ModFOLDdock improves ranking agreement

The reliability, however, was not maintained for all multimers. As shown by the plots in section 3.1.2, both pTM and plDDT showed variability for models of similar quality with pTM showing a tendency for underestimation for higher quality models and both scores showing overestimation for some lower quality models which was more pronounced for plDDT. This variability affected ranking accuracy, with both pTM and plDDT showing a lower association with observed score ranking than was seen for monomers. Of the two scores the association was less strong for plDDT-assigned ranks.

ModFOLDdock, which did not show over-prediction to the same degree, was able to improve upon the rank agreement of plDDT although there was insufficient evidence to draw the same conclusion for pTM. Nevertheless, there remains some unreliability in the ability of pTM and plDDT to differentiate between some high and low quality multimer models created by regular modelling and ModFOLDdock scores represent a more reliable method for ranking multimer models.

### Greater variability of AF2 predicted scores is seen if custom template recycling is used

Finally, convincing evidence is presented in Supplementary section S3.4 that using the custom template option to recycle models through the AlphaFold2 algorithm results in a much greater variability in predicted scores for both tertiary structures and multimers and that the variability is more extreme for multimer models. This provides cautionary evidence that the use of AF2 and AF2-Multimer outside of their intended end-to-end operation could result in mis-scoring and mis-ranking of models.

### Independent MQA is essential for multimer models

For AF2-Multimer modelling, whether straight forward or via custom template recycling, ModFOLDdock should also be used as an independent MQA program. This not only offers an alternative opinion on quality and enables models from different software to be objectively compared but also appears less prone to variation in prediction across the model quality range.

## Supporting information

Supplementary section

## References

Adiyaman, R., Edmunds, N. S., Genc, A. G., Alharbi, S. M. A. & Mcguffin, L. J. 2023. Improvement of protein tertiary and quaternary structure predictions using the ReFOLD refinement method and the AlphaFold2 recycling process. Bioinform Adv, 3, vbad078.

Evans, R., O’neill, M., Pritzel, A., Antropova, N., Senior, A., Green, T., Žídek, A., Bates, R., Blackwell, S., Yim, J., Ronneberger, O., Bodenstein, S., Zielinski, M., Bridgland, A., Potapenko, A., Cowie, A., Tunyasuvunakool, K., Jain, R., Clancy, E., Kohli, P., Jumper, J. & Hassabis, D. 2022. Protein complex prediction with AlphaFold-Multimer. bioRxiv, 2021.10.04.463034.

Jeffrey Skolnick, Statesmu Gao, Hongyi Zhou & Singh, S. 2021. AlphaFold 2: Why It Works and Its Implications for Understanding the Relationships of Protein Sequence, Structure, and Function. J Chem Inf Model, 61.

Jumper, J., Evans, R., Pritzel, A., Green, T., Figurnov, M., Ronneberger, O., Tunyasuvunakool, K., Bates, R., Zidek, A., Potapenko, A., Bridgland, A., Meyer, C., Kohl, S. a. A., Ballard, A. J., Cowie, A., Romera-Paredes, B., Nikolov, S., Jain, R., Adler, J., Back, T., Petersen, S., Reiman, D., Clancy, E., Zielinski, M., Steinegger, M., Pacholska, M., Berghammer, T., Bodenstein, S., Silver, D., Vinyals, O., Senior, A. W., Kavukcuoglu, K., Kohli, P. & Hassabis, D. 2021. Highly accurate protein structure prediction with AlphaFold. Nature, 596, 583–589.

Mariani, V., Biasini, M., Barbato, A. & Schwede, T. 2013. lDDT: a local superposition-free score for comparing protein structures and models using distance difference tests. Bioinformatics, 29, 2722–8.

Mcguffin, L. J., Edmunds, N. S., Genc, A. G., Alharbi, S. M. A., Salehe, B. R. & Adiyaman, R. 2023. Prediction of protein structures, functions and interactions using the IntFOLD7, MultiFOLD and ModFOLDdock servers. Nucleic Acids Res, 51, W274–W280.

Mirdita, M., Schutze, K., Moriwaki, Y., Heo, L., Ovchinnikov, S. & Steinegger, M. 2022. ColabFold: making protein folding accessible to all. Nat Methods, 19, 679–682.

Richard Evans, M. O. N., Alexander Pritzel, Natasha Antropova, Andrew Senior, Tim Green, Augustin Žídek, Russ Bates, Sam Blackwell, Jason Yim, Olaf Ronneberger, Sebastian Bodenstein, Michal Zielinski, Alex Bridgland, Anna Potapenko, Andrew Cowie, Kathryn Tunyasuvunakool, Rishub Jain, Ellen Clancy, Pushmeet Kohli, John Jumper, Demis Hassabis 2021. Protein complex prediction with AlphaFold-Multimer. bioRxiv.

Roney, J. P. & Ovchinnikov, S. 2022. State-of-the-Art Estimation of Protein Model Accuracy Using AlphaFold. Phys Rev Lett, 129, 238101.

Sergey Ovchinnikov, Martin Steinegger & Mirdita, M. 2022. Benchmarking ColabFold in CASP15. CASP15 Abstracts, 50.

Shao, C., Bittrich, S., Wang, S. & Burley, S. K. 2022. Assessing PDB macromolecular crystal structure confidence at the individual amino acid residue level. Structure, 30, 1385–1394.

Stroe, O. 2021. Pfam releases structures for every protein family [Online]. Available: https://www.ebi.ac.uk/about/news/announcements/Pfam-protein-structures/ [Accessed].

Terwilliger 2022. Improving AlphaFold modeling using implicit information from experimental density maps. BioRxiv.

Tunyasuvunakool, K., Adler, J., Wu, Z., Green, T., Zielinski, M., Zidek, A., Bridgland, A., Cowie, A., Meyer, C., Laydon, A., Velankar, S., Kleywegt, G. J., Bateman, A., Evans, R., Pritzel, A., Figurnov, M., Ronneberger, O., Bates, R., Kohl, S. a. A., Potapenko, A., Ballard, A. J., Romera-Paredes, B., Nikolov, S., Jain, R., Clancy, E., Reiman, D., Petersen, S., Senior, A. W., Kavukcuoglu, K., Birney, E., Kohli, P., Jumper, J. & Hassabis, D. 2021. Highly accurate protein structure prediction for the human proteome. Nature, 596, 590–596.

Varadi, M., Anyango, S., Deshpande, M., Nair, S., Natassia, C., Yordanova, G., Yuan, D., Stroe, O., Wood, G., Laydon, A., Zidek, A., Green, T., Tunyasuvunakool, K., Petersen, S., Jumper, J., Clancy, E., Green, R., Vora, A., Lutfi, M., Figurnov, M., Cowie, A., Hobbs, N., Kohli, P., Kleywegt, G., Birney, E., Hassabis, D. & Velankar, S. 2022. AlphaFold Protein Structure Database: massively expanding the structural coverage of protein-sequence space with high-accuracy models. Nucleic Acids Res, 50, D439–D444.

Wallner, B. 2023. AFsample: Improving Multimer Prediction with AlphaFold using Aggressive Sampling. bioRxiv, 2022.12.20.521205.

Yuma Takei & Ishida, T. 2022. How to select the best model from AlphaFold2 structures? bioRxiv.

Zhang, Y. & Skolnick, J. 2004. Scoring function for automated assessment of protein structure template quality. Proteins, 57, 702–10.

