## Supplementary section for "Benchmarking of AlphaFold2 accuracy self-estimates as empirical quality measures and model ranking indicators and their comparison with independent model quality assessment programs": BioRXiv_Supplementary.pdf

**S1.1 Evans' description of ranking used in later versions of AlphaFold2 and ColabFold.**
Ranking was calculated as a weighted combination of pTM and interface ipTM, calculated as (0.8 x ipTM + 0.2 x pTM). ColabFold v1.5.0 (Jan-2022 onwards) used the weighted ipTM-pTM score to rank multimers when using the AlphaFold2_mmseqs2, AlphaFold2_batch and colabfold_batch variants.

**S2.2 Population A (CASP15 monomers).**
McGuffin group's submissions for CASP15 regular tertiary structure for 26 targets: T1104, T1112, T1120, T1122, T1125, T1130, T1131, T1133, T1139, T1145, T1146, T1147, T1150, T1154, T1155, T1158, T1159, T1162, T1163, T1175, T1177, T1180, T1182, T1183, T1188 & T1194.

**S2.3 Population B (CASP15 multimers).**
Scores for rank 1 to 5 models were collected for all multimer models for which data were available, resulting in 395 individual models across 41 targets (the ColabFold group submitted no models for three targets making a total of 38); H1106, H1111, H1114, H1129, H1134, H1135, H1137 (MultiFOLD only), H1140, H1141, H1142, H1143, H1144, H1151, H1157, H1166, H1167, H1168, H1171, H1172, H1185, T1109, T1110, T1113, T1115 (MultiFOLD only), T1121, T1123, T1124, T1127, T1132, T1153, T1160, T1161, T1170, T1173, T1174, T1176, T1178, T1179, T1181, T1187 and T1192 (MultiFOLD only).

**S2.4 Population C (recycled monomers).**
The AlphaFold2 Rank 1 models were downloaded from the CASP14 website for the following 20 CASP14 FM targets: T1027, T1029, T1031, T1033, T1037, T1039, T1040, T1041, T1042, T1043, T1047s1, T1047s2, T1055, T1058, T1064, T1074, T1090, T1093, T1094, T1096. Again, as described in section S2.3, observed quality assessment scores were generated using the downloadable versions of TM-score and lDDT score. To affect the recycling, model PDB files were converted to mmCIF format using the RSCB PDB MAXIT suite of programs (https://mmcif.pdbj.org/converter). These were then submitted to the Google Colaboratory hosted ColabFold (release 3, v1.3.0 [4-Mar-2022]) as custom templates along with their respective amino acid sequences. ColabFold was run twice per model (both MSA and single-sequence modes), and, within each mode, the model was submitted four times for 1, 3, 6 and 12 recycles. ColabFold settings used were: Template_mode: custom; msa_mode: MMseqs2 (UniRef+Environmental) OR single sequence; pair_mode: unpaired+paired; model-type: auto; num_recycles: 1, 3, 6, 12 (selecting "auto" from the model type defaulted to the original pre-CASP14 AF2 model). Amber relaxation was not enabled. Models created for each ColabFold run were collected along with their predicted pTM and plDDT scores and then rescored with TM-score and lDDT as described above.

The same logic was employed for nonAF2 CASP14 models. These were selected from the same 20 FM targets for the next five best-ranked groups beneath AlphaFold2 at CASP14. These were Baker (473), Baker-experimental (403), Feig-R2 (480), Zhang (129) and tFold_human (009). To ensure consistency in terms of globular fold similarity, only models with a TM-score >=0.45 were selected and this resulted in a total of 47 individual models. The full list of models used is:
T1029TS009_1-D1, T1031TS009_1-D1, T1033TS009_1-D1, T1037TS009_1-D1, T1041TS009_1-D1, T1042TS009_1-D1, T1043TS009_1-D1, T1049TS009_1-D1, T1090TS009_1-D1, T1031TS129_1-D1, T1037TS129_1-D1, T1040TS129_1-D1, T1041TS129_1-D1, T1042TS129_1-D1, T1049TS129_1-D1, T1074TS129_1-D1,

T1090TS129_1-D1, T1096TS129_1, T1027TS403_1-D1, T1031TS403_1-D1, T1033TS403_1-D1, T1037TS403_1-D1, T1039TS403_1-D1, T1041TS403_1-D1, T1042TS403_1-D1, T1043TS403_1-D1, T1049TS403_1-D1, T1090TS403_1-D1, T1096TS403_1, T1031TS473_1-D1, T1033TS473_1-D1, T1037TS473_1-D1, T1039TS473_1-D1, T1041TS473_1-D1, T1042TS473_1-D1, T1043TS473_1-D1, T1049TS473_1-D1, T1074TS473_1-D1, T1090TS473_1-D1, T1031TS480_1-D1, T1037TS480_1-D1, T1041TS480_1-D1, T1042TS480_1-D1, T1049TS480_1-D1, T1074TS480_1-D1, T1090TS480_1-D1, T1096TS480_1.

Models were downloaded from the CASP14 website, scored with TM-score and lDDT and modelled with the MSA option in the same way as described for AF2 models. Single sequence modelling was carried out using release v1.3.0 of Localcolabfold (Mirdita *et al.*, 2022) installed on our own server, to overcome the Google Colaboratory GPU restrictions in the time available. The equivalent Localcolabfold settings were used: msa-mode: single_sequence; model-type: auto; rank: plddt; pair-mode: unpaired+paired; templates: --custom-template-path. The resulting rank 1-5 models were collected along with their plDDT and pTM scores and rescored against the native structure to produce a set of observed lDDT and TM-scores.

**S2.5 Population D (recycled multimers)**

Some of multimer targets were too large to recycle through AF2-Multimer (training was limited to models up to 1536 residues and the algorithm can experience memory issues with models of more than a few thousand residues (Bryant *et al.*, 2022) and therefore the targets used in this set were limited by size to: H1045, H1065, H1072, T1032, T1054, T1070, T1073, T1078, T1083, T1084. Again, top-ranked models were used as the custom templates and were subjected to recycling (1, 3, 6 and 12) using ColabFold (MSA and SS modes) in the same way as described for the monomer structures above. The resulting 50 rank 1-5 models were then collected along with their plDDT and pTM scores. Observed scores were obtained by assessing each model against their relevant native structures using the OpenStructure and MM-Align (Mukherjee and Zhang, 2009) programs to obtain observed oligo_lDDT and TM-scores respectively.

**S2.6 Handling of contingency table data and Multimer pTM scores and the procedure for model ranking.**

Multimer models created by AlphaFold2 variants are, by default, ranked by pTM rather than plDDT. As stated in the introduction there is a slight difference in the calculation of the pTM-based ranking between versions of ColabFold. In AF2-Multimer and in later versions of ColabFold (v1.5.0) ranking is calculated based on a ratio of 0.8 * ipTM + 0.2 * pTM (Richard Evans, 2021), whereas in earlier versions, ranking is calculated on pTM score alone. As some multimer models in this population were created with ColabFold v1.3 and some with v1.5 there was potentially heterogeneous ranking across the model population, and it was necessary to allow for this when comparing ranks. To this end, multimer models were routinely re-ranked by pTM score before comparison with observed rankings. The procedure for deriving model ranks in R consisted of ranking each individual set of 5 related models, i.e., models output from a single run of AlphaFold2 modelling, using the statement *rank <score>, ties.method ="random"* where <score> can be replaced with any of the predicted or observed scores as necessary. This was applied to observed score ranking but also to ranking by pTM for the reasons explained above. In this way any differences in the way the AlphaFold2 algorithm originally ranked the data were negated and the ranks were assigned uniformly across all populations.

Multi-factor contingency tables to display ranking comparisons were created in R using the caret package with the confusionMatrix() command and four further statistical measures were used to assess relatedness. Sensitivity, specificity, precision, and accuracy were calculated for individual rank classes (1, 2, 3, 4 and 5) and, to construct meaningful comparisons between the contingency tables, macro-averaged versions of these statistics were calculated as mean values across all categories for each table.

Fisher's exact test is often used for smaller sample sizes (single contingency table cells of less than 5) or where independence of observations cannot be guaranteed and, while the concept of independence holds for the assignment of ranks based on predicted and observed scores, some tables do have low figures in individual cells. As regards multi-contingency tables (larger than 2x2), no clear distinction between the two tests could be found other than Chi-squared may run into problems with very sparse data and Fisher's can become computationally intensive for large tables.

As ranking data is categorical, it is possible to assess the association between the predicted model ranks and observed model ranks using the Chi-squared and Fisher's exact tests, where P-values $<0.05$ would suggest relatedness between distributions. It was decided that both tests would be run as a check for each other, i.e., agreement between the two tests would confer confidence in the result. A Monte Carlo resampling method (simulate.p.value) with default simulations of 2000 was used for the Fisher's exact test to allow a more robust estimate of the p-value and prevent any computational overheads which can occur when this test is applied to larger contingency tables (Crawley, 2015). Analysis was performed using R version 3.6.3 running in R-studio.

#### S3.4 Hypothesis 4. Is the accuracy of predicted scores affected by custom template recycling?

To answer this question data is presented from the four model populations which underwent custom template recycling. For monomers this is Population A2 (CASP15 round 2 monomers) and Population C (recycled monomers), for multimers it is Population B2 (CASP15 MultiFOLD group multimers) and Population D (recycled multimers). It would be logical to start with the data for populations A2 and B2 because these two groups can be directly compared to their unrecycled counterparts, i.e. Population A2, the CASP15 round 2 monomers (recycled) can be directly compared with the Population A1 CASP15 round 1 monomers (unrecycled) which were discussed in section 3.1.1 and Population B2, the MultiFOLD group multimers (recycled) can be directly compared to the Population B1 ColabFold group multimers (unrecycled) which were discussed in section 3.1.2. Populations C and D have no direct comparisons and so will be discussed last to provide support of the population A and B data.

### S3.4.1 Population A2 (CASP15 round 2 monomers).

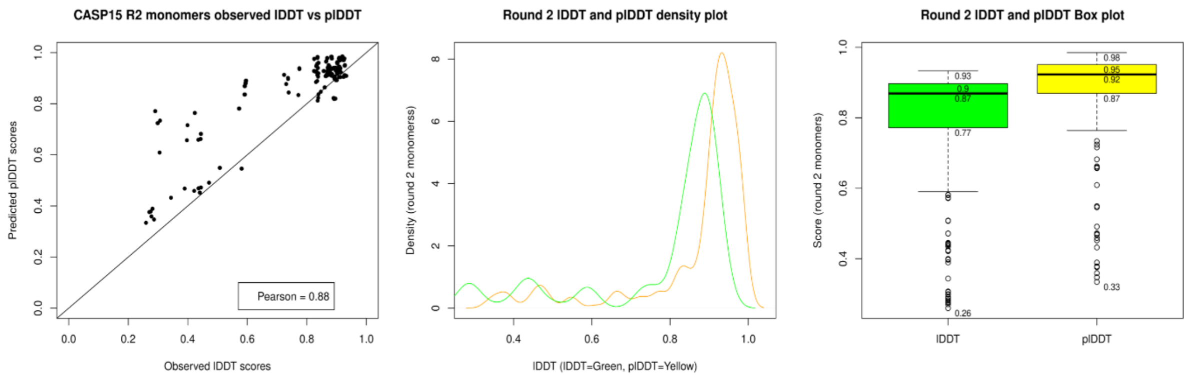


**Figure S11. Plots for plDDT versus observed lDDT for Population A2 (CASP15 round 2 monomers)**. A scatter plot (left), density plot (middle) and boxplot (right). For all plots plDDT has been rescaled to fit the 0-1 lDDT range.

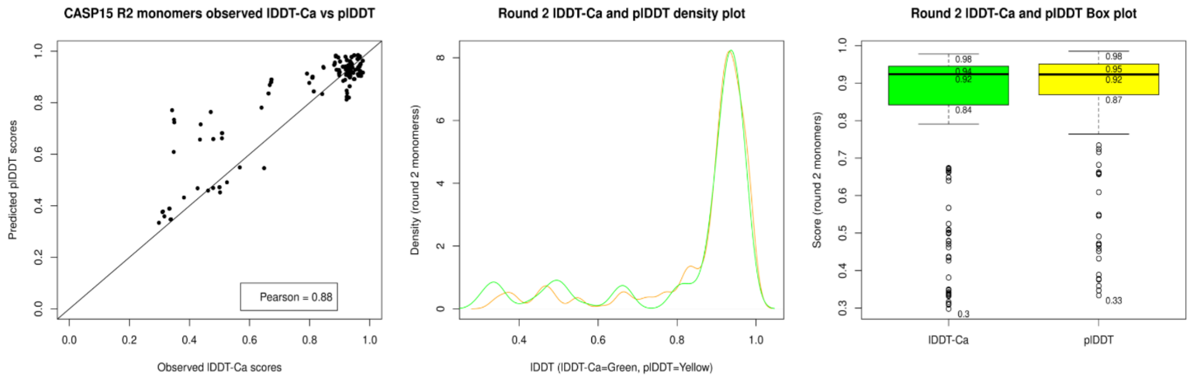


**Figure S12. Plots for plDDT versus observed lDDT-Cα for Population A2 (CASP15 round 2 monomers)**. A scatter plot (left), density plot (middle) and boxplot (right). For all plots plDDT has been rescaled to fit the 0-1 lDDT range.

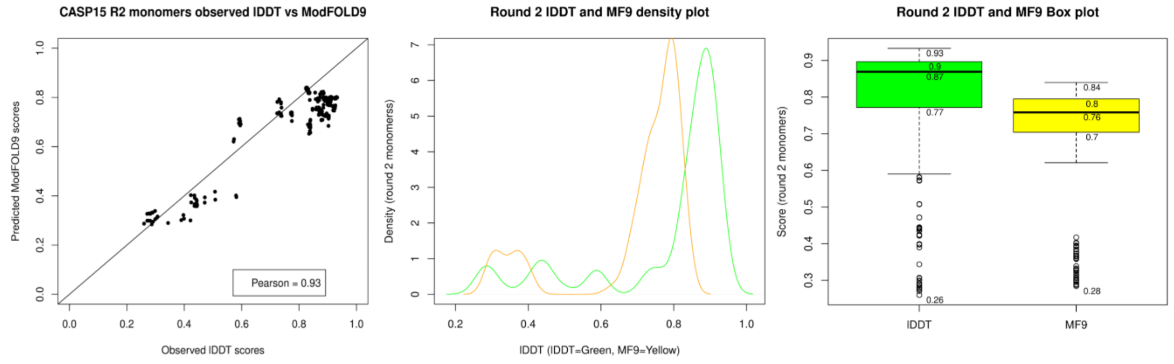


**Figure S13. Equivalent plots of ModFOLD9 score versus observed lDDT for Population A2 (CASP15 round 2 monomers).** A scatter plot (left), density plot (middle) and boxplot (right). For all plots plDDT has been rescaled to fit the 0-1 lDDT range.

Comparing the data from Figure S11 directly with that for the round 1 monomers presented in Figure 2, it is apparent that there is a wider spread of data in the scatter plot in Figure S11 with an increase in plDDT scores, which are reflected in the density plot and the boxplot. Figure S12, for lDDT-Cα scores, shows a similar spread in the scatter plot but accompanied by a less noticeable difference between the plDDT and lDDT-Cα distributions in the density and boxplot. Wilcoxon signed rank tests for significance in Table S8 (below), however, reveal that the difference between the plDDT and lDDT-Cα score is significant as is the difference between the round 2 monomer plDDT scores and their round 1 counterparts.

**Table S8. Wilcoxon statistics for Population A2 round 2 monomers.** (significant figures in bold).

| *Row* | *Scores compared* | *Independence and distribution symmetry* | *p-value* |
|---|---|---|---|
| *1* | *R2 plDDT versus lDDT-Cα* | Paired; 2-sided test | **0.0001** |
| *2* | *R2 plDDT versus lDDT-Cα* | Paired; 1-sided test, plDDT > lDDT-Cα | **$5.83 \times 10^{-5}$** |
| *3* | *R2 plDDT versus R1 plDDT* | Unpaired; 2-sided | **$1.293 \times 10^{-9}$** |
| *4* | *R2 plDDT versus R1 plDDT* | Unpaired; 1-sided, R2 > R1 | **$6.465 \times 10^{-10}$** |
| *5* | *R2 lDDT-Cα versus R1 lDDT-Cα* | Unpaired; 2-sided | 0.1255 |

Table S8, row 1, shows that according to a paired 2-sided Wilcoxon test there is a significant difference between plDDT and lDDT-Cα observed scores for round 2 monomers and, further to this, the results of a paired 1-sided test in row 2 show that that plDDT scores are significantly higher. These findings agree with the scatter plots in Figures S11 and S12 showing over-prediction in mid-quality models which was not present in the round 1 data. Notably, the over-prediction is also absent from the equivalent round 2 ModFOLD9 scatter plot shown in Figure S13. This is good evidence that overprediction of plDDT occurs in monomer models with custom template recycling.

To further test this, a 2-sided Wilcoxon test was used to directly compare round 1 and round 2 monomer plDDT scores (row 3) and this showed a significant difference between the two scores, evidenced by a p-value of $1.293 \times 10^{-9}$. Further, it was established that the round 2 monomer scores were significantly higher than those for round 1, evidenced by a p-value of $6.465 \times 10^{-10}$ from the 1-sided test in row 4. Importantly, there was no such difference between the equivalent round 1 and 2 monomer observed lDDT-Cα scores as shown by the p-value of 0.1255 (row 5 of the table) meaning that round 1 and 2 monomer models were not significantly different in quality.

It is therefore reasonable to conclude that these prediction errors have been introduced by custom template recycling and, for hypothesis 4 in respect to monomer models, the alternative hypothesis can be accepted, i.e., *AF2 predicted scores following custom template modelling show greater variation than scores from regular modelling, when compared to equivalent observed scores.*

### S3.4.2 Population B2 (CASP15 MultiFOLD multimers).

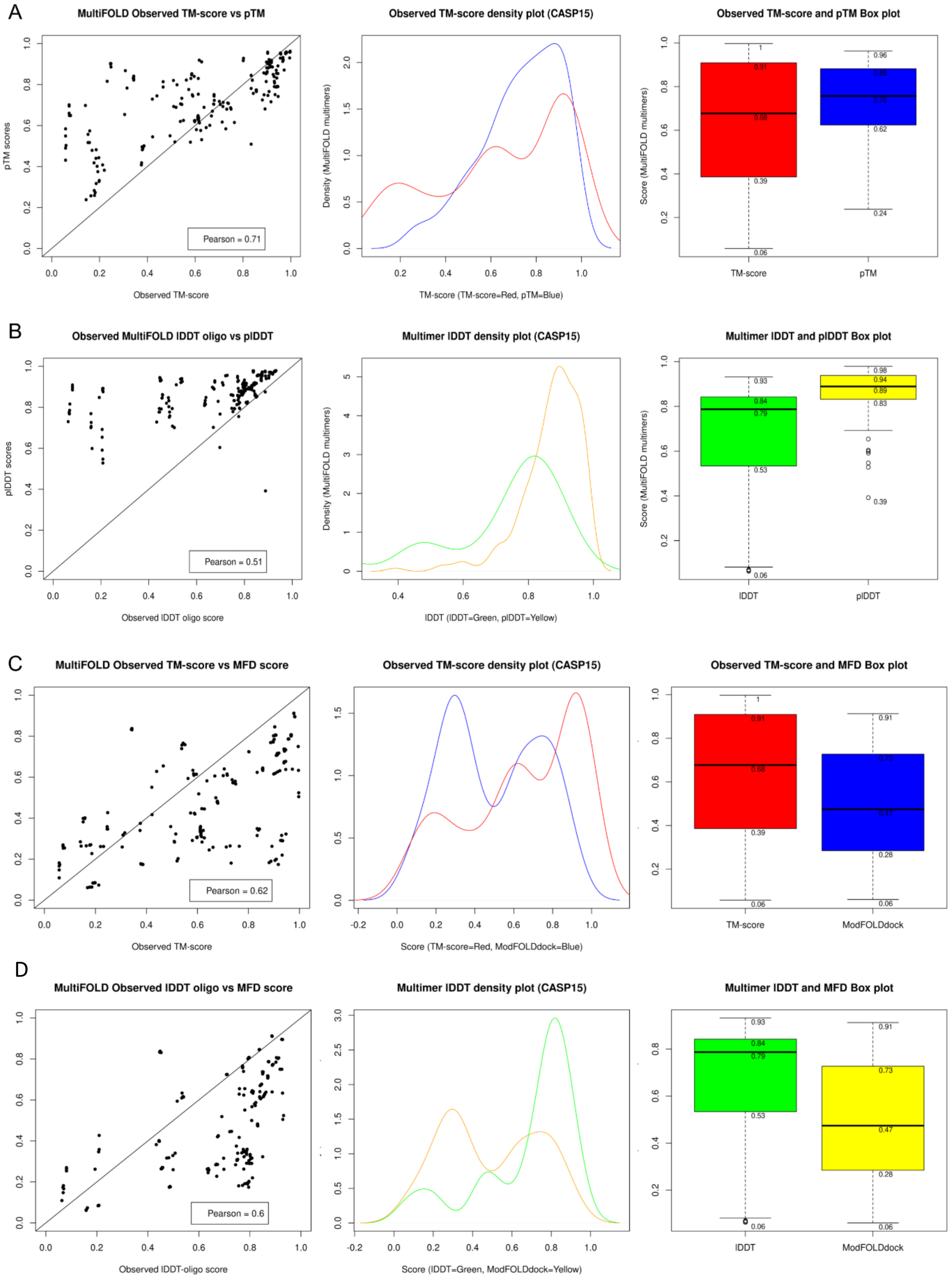


**Figure S14. Plots for Population B2 (MultiFOLD multimers). scatter plots (left), density plots (middle) and box plots (right) for.** Panel A: pTM versus observed TM-score. Panel B: plDDT versus observed CASP lDDT-oligo. Panel C: Comparison plots for ModFOLDdock score versus TM-score, Panel D Comparison plots for ModFOLDdock versus lDDT-olio.

The plots in Figure S14, panels A and B, can be directly compared to Figures 4 and 5 for ColabFold multimers in section 3.1.2. Considering the plots in panel A for TM-scores, the spread of points in the scatter plot is again noticeably greater than that shown in Figure 4. Further, although the mean observed TM-score in the boxplots reduces from 0.745 (Figure 4) to 0.68 across the two populations, the equivalent mean pTM rises from 0.72 to 0.76. Secondly, considering panel B for lDDT scores in a similar way, the scatter plot again shows an increase in the spread of data compared to its equivalent in Figure 5. and there is also a marked shift to the right in plDDT when comparing the density plots, and a corresponding increase in mean plDDT score shown in the boxplot. These changes suggest a similar overprediction to that seen for monomers is also occurring for multimers which have been subject to custom template recycling. For comparison, the scatter plots in panel C showing ModFOLDdock scores versus both observed TM-score and lDDT-oligo scores for the same population, show little evidence of sustained overprediction. If anything, ModFOLDdock appears to suffer from a tendency for under-prediction of these models.

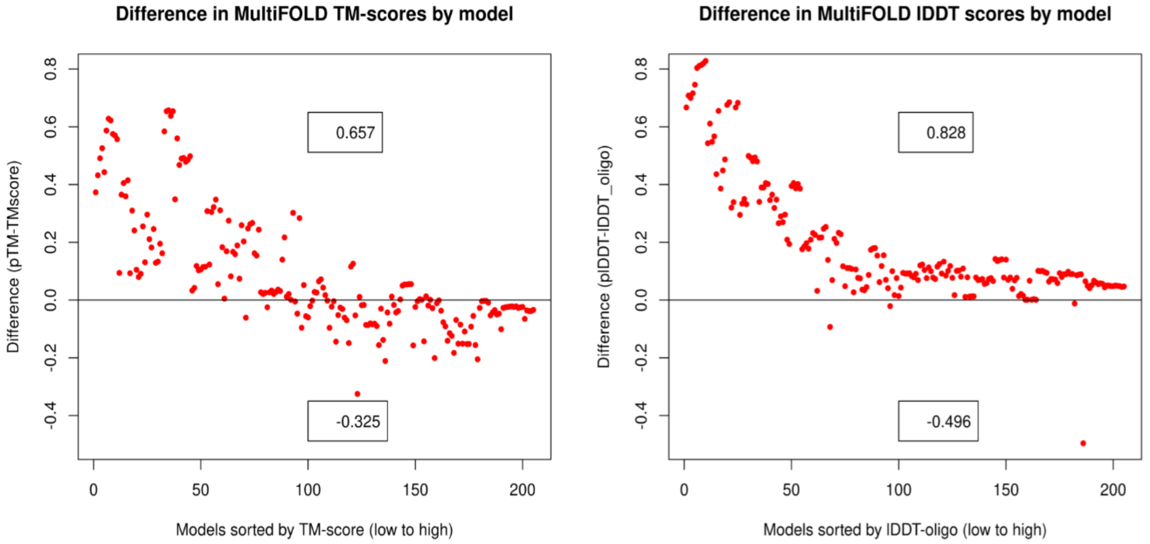


**Figure S15. Plots to show variation between predicted and observed scores for Population B2 (MultiFOLD multimers).** Left, pTM versus TM-score and right, plDDT versus lDDT-oligo. Plots are equivalent to those in Figure 6 for ColabFold multimers.

The relationships suggested in Figure S14, panels A and B, are more clearly shown by the two variation plots in Figure S15. In agreement with Figure 6, both plots show overprediction of scores for lower quality models with maximum and minimum differences of +0.657 and -0.325 respectively for pTM score and a maximum difference of +0.828 for plDDT score. Although the maximum and minimum deviation in the data for pTM score are almost identical to those from Figure 6, the maximum deviation in plDDT scores has increased from 0.747 to 0.828. Also, upon visual comparison of the two pairs of plots it is clear that the number models in the over-predicted regions in Figure S15 has increased over those in Figure 6 despite a similar number of models (205 and 190 respectively). Wilcoxon signed rank tests were again used to quantify these differences in terms of significance and the results are presented in Table S9 below.

**Table S9. Wilcoxon tests for Population B2 MultiFOLD multimers and Population B1 ColabFold multimers.** (significant figures in bold).

| *Row* | *Scores compared* | *Independence and distribution symmetry* | *p-value* |
|---|---|---|---|
| *1* | *MultiFOLD plDDT versus lDDT-oligo* | Paired; 1-sided test, plDDT > lDDT-oligo | **$2.20\times10^{-16}$** |

| | | | |
|---|---|---|---|
| *2* | *MultiFOLD pTM versus TM-score* | Paired; 1-sided test, pTM > TM-score | **$1.46\text{x}10^{-5}$** |
| *3* | *MultiFOLD versus ColabFold plDDT* | Unpaired; 1-sided, MultiFOLD > ColabFold | **$7.193\text{x}10^{-8}$** |
| *4* | *MultiFOLD versus ColabFold lDDT-oligo* | Unpaired; 2-sided. | 0.283 |
| *5* | *MultiFOLD versus ColabFold pTM* | Unpaired; 1-sided; MultiFOLD > ColabFold | **0.014** |
| *6* | *MultiFOLD versus ColabFold TM-score* | Unpaired; 2-sided. | 0.252 |

Table S9, rows 1 and 2 confirm that both predicted plDDT and pTM scores are significantly greater than their observed counterparts (lDDT and TM-scores) for MultiFOLD multimers as evidenced by p-values of $2.20\text{x}10^{-16}$ for plDDT versus lDDT-oligo and $1.46\text{x}10^{-5}$ for pTM versus TM-score. Furthermore, there is confirmation that MultiFOLD plDDT (row 3) and pTM (row 5) scores are significantly greater than the equivalent predicted scores for ColabFold multimers but, importantly, there is no significant difference between the equivalent two sets of observed scores (row 4 for lDDT-oligo and row 6 for TM-score). This again shows that, for a similar set of models based on the same CASP targets, both sets of observed scores are similar but both predicted pTM and plDDT scores are significantly different and are higher in both cases for the group subject to custom template recycling.

Therefore, with respect to multimers, the alternative hypothesis must again be accepted, i.e., *AF2 predicted scores following custom template modelling show greater variation than scores from regular modelling, when compared to equivalent observed scores.*

#### S3.4.3 Population C (recycled monomers).

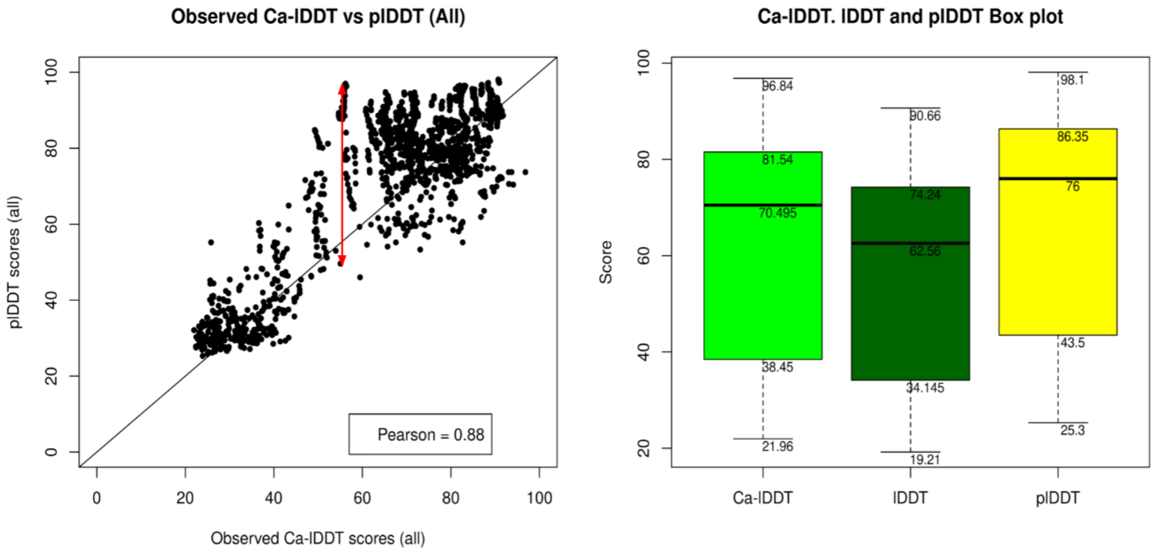


**Figure S16. Plots for plDDT versus observed lDDT-Cα for population C (recycled monomers).** Left, a scatter plot showing the spread of data and right, a boxplot comparing the distribution of lDDT-Cα, lDDT and plDDT scores for the same population. lDDT and lDDT-Cα have been rescaled to the 0-100 range.

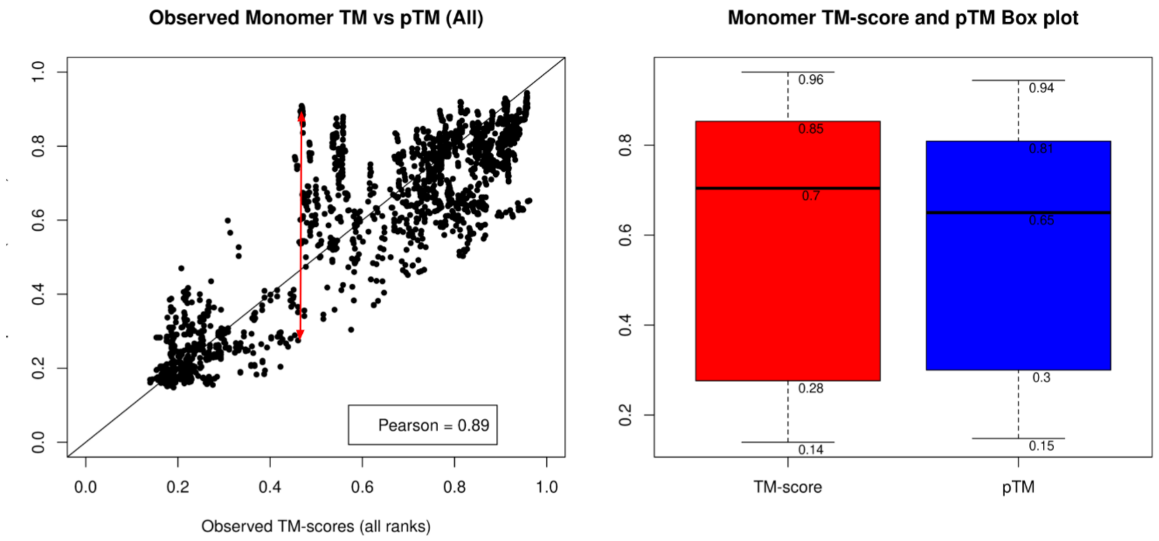


**Figure S17. Plots for pTM versus observed TM-score for population C (recycled monomers).** Left, a scatter plot showing the spread of data and right, a boxplot for both scores from the same population.

The scatter plots in Figures S16 and S17 show Pearson correlation coefficients of 0.87 and 0.89 respectively between predicted and observed scores. Although these correlations appear very respectable, both plots show a pronounced spread in the data with a high proportion of outliers. The red bars on each scatter plot show the potential degree of variation in predicted scores for models with similar observed scores. For an observed score of approximately 0.5, predicted plDDT scores range from approximately 0.5 to 0.9 (Figure S16) and pTM scores range from approximately 0.3 to 0.9 (Figure S17).

These results strongly support the hypothesis that using custom template recycling appears to produce a much higher degree of variability both plDDT and pTM scores.

#### S3.4.4 Population D (recycled multimers).

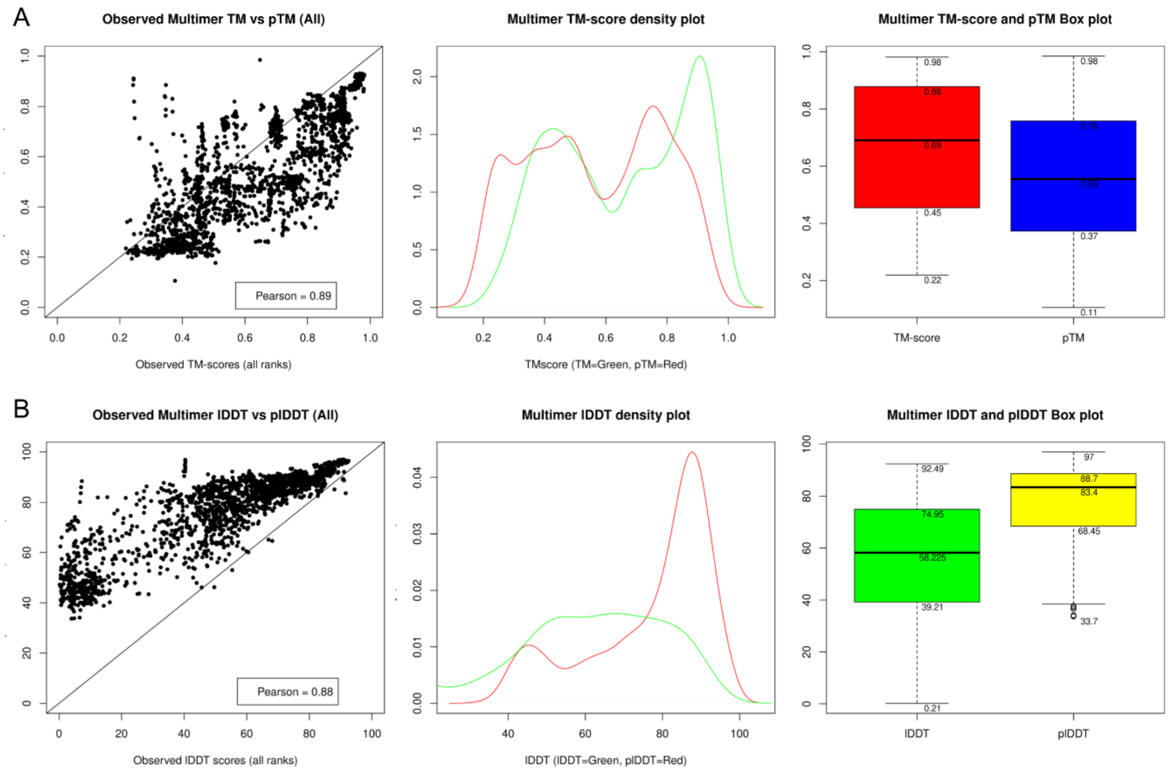


**Figure S18. Plots for Population D (Recycled multimers). Scatter plots (left), density plots (middle) and box plots (right).** Panel A: pTM versus observed TM-score and panel B: plDDT versus observed CASP lDDT-oligo.

Figure S18 shows a similar spread of data to that seen in Figures S16 and S17. Panel A again shows a tendency for multimer pTM over and under-prediction meaning a high variation in predicted pTM score for models with similar observed scores. In panel B, all three plots demonstrate a high tendency for plDDT over-prediction and again, this is more pronounced for mid to lower quality models.

As both population C and D were subject to up to 12 recycles and were entirely created via custom template recycling, these results support the hypotheses drawn above for population A and B, that using custom template recycling produces a higher degree of variability in AlphaFold2 predicted scores for both monomer and multimer models and that this effect is more pronounced for multimers.
